# Whole-genome duplication in an algal symbiont bolsters coral heat tolerance

**DOI:** 10.1101/2022.04.10.487810

**Authors:** Katherine E. Dougan, Anthony J. Bellantuono, Tim Kahlke, Raffaela M. Abbriano, Yibi Chen, Sarah Shah, Camila Granados-Cifuentes, Madeleine J. H. van Oppen, Debashish Bhattacharya, David J. Suggett, Mauricio Rodriguez-Lanetty, Cheong Xin Chan

## Abstract

The algal endosymbiont *Durusdinium trenchii* enhances the resilience of coral reefs under thermal stress^1,2^. As an endosymbiont, *D. trenchii* is generally expected to have a reduced genome compared to its free-living relatives, due in part to the lack of selective pressure for maintaining redundant gene functions in a stable intracellular environment within the host^3^. However, *D. trenchii* can live freely or in endosymbiosis, and the analysis of genetic markers^4^ suggests that this species has undergone whole-genome duplication (WGD). Here we present genome assemblies for two *D. trenchii* isolates, confirm WGD in these taxa, and examine how selection has shaped the duplicated genome regions. We assess how the competing free-living versus endosymbiotic lifestyles of *D. trenchii* have contributed to the retention and divergence of duplicated genes, and how these processes have enhanced thermotolerance of corals hosting these symbionts. We find that lifestyle is the driver of post-WGD evolution in *D. trenchii*, with the free-living phase being most important, followed by endosymbiosis. Adaptations to both lifestyles collectively result in increased cellular fitness for *D. trenchii*, which provides enhanced thermal stress protection to the host coral. Beyond corals, this polyploid alga is a valuable model for understanding how genome-wide selective forces act to balance the often, divergent constraints imposed by competing lifestyles.

## Main text

Uncovering the foundations of biotic interactions, particularly symbiosis, remains a central goal for research, given that virtually no organism lives in isolation. Coral reefs are marine biodiversity hotspots that are founded upon symbioses involving dinoflagellate algae in the Family Symbiodiniaceae^5^. These symbionts are the “solar power plants” of reefs, providing photosynthetically fixed carbon and other metabolites to the coral host^6,7^. Breakdown of the coral-dinoflagellate symbiosis (i.e. coral bleaching), often due to ocean warming, puts corals at risk of starvation, disease, and eventual death. Symbiodiniaceae microalgae are diverse, with at least 15 clades including 11 named genera^5,8–10^, encompassing a broad spectrum of symbiotic associations and host-specificity. Most of these taxa are facultative symbionts (i.e. they can live freely or in symbiosis), although exclusively symbiotic or free-living species also exist^5^. Genomes of Symbiodiniaceae are believed to reflect the diversification and specialization of these taxa to inhabit distinct ecological niches^3,11^. The genomes of symbionts, due to spatial confinement, are predicted to undergo structural rearrangements, streamlining, and rapid genetic drift (e.g. pseudogenization)^3^. These traits are present in symbiotic Symbiodiniaceae^11^.

Whole-genome duplication (WGD) is an evolutionary mechanism for generating functional novelty and genomic innovation^12,13^, and can occur due to errors in meiosis, i.e. via autopolyploidy. Following WGD, the evolutionary trajectory of duplicated sequence regions generally proceeds from large-scale purging, temporary retention and/or divergence, to fixation^14^. WGD-derived duplicated genes (i.e. *ohnologs*^15,16^) that are retained can provide a selective advantage and enhance fitness through increased gene dosage, specialization in function, and/or the acquisition of novel functions^14^.

WGD has been described in free-living unicellular eukaryotes such as yeast^17–19^, ciliates^20,21^, and diatoms^22,23^, but not in symbiotic species. Evidence of WGD is absent in the Symbiodiniaceae, except for the genus *Durusdinium*, as observed in microsatellite sequence data^4^. This genus includes the thermotolerant species *Durusdinium trenchii* (Fig. 1a), a facultative symbiont that confers heat-tolerance to corals, thereby enhancing holobiont resilience under thermal stress^24^. We hypothesize that WGD played a critical role in enhancing heat-tolerance in this species. Specifically, the facultative lifestyle (i.e. free-living or symbiotic) of *D. trenchii* favoured fixation of WGD during the free-living phase as an adaptation to fluctuating environmental conditions, with the expanded gene inventory being further modified by the coral symbiosis. To test this “dual lifestyle” hypothesis, we generated *de novo* genome assemblies from two isolates of *D. trenchii* and analysed their evolutionary trajectories. Predictions of our hypothesis were tested against the null model of a single, free-living, lifestyle for this species. Based on gene expression profiles, we elucidate how the facultative lifestyle has contributed to the fate of ohnologs in these microalgae, and how natural selection acting on gene families has increased thermotolerance of corals hosting *D. trenchii* symbionts. These data provide strong evidence for the dual lifestyle hypothesis as a driver of post-WGD genome evolution.

**Figure 1.**
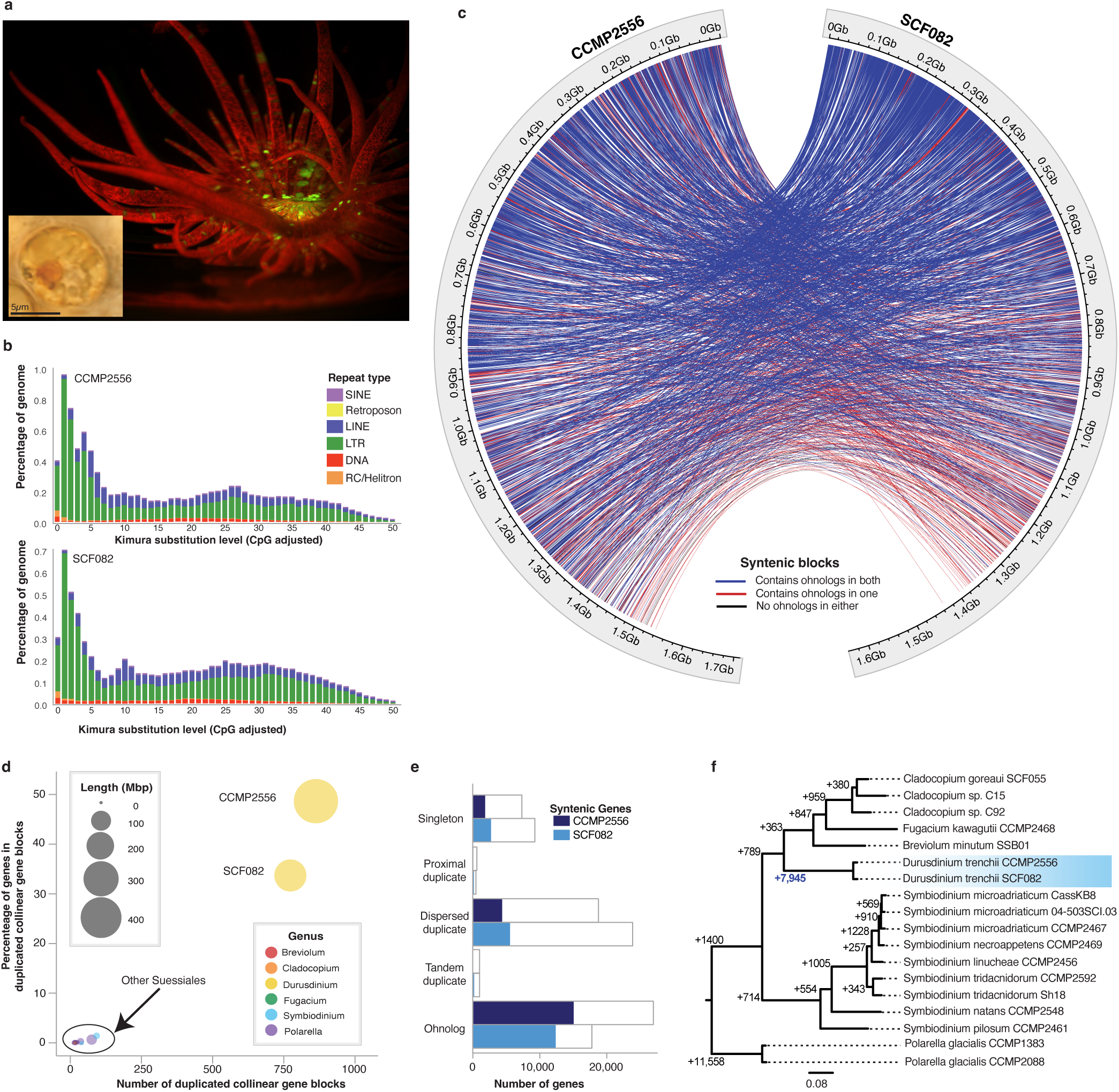
WGD in a facultative coral endosymbiont. (**a**) Microscopic images of a free-living *D. trenchii* cell and a *Exaiptasia pallida* anemone hosting *D. trenchii* under fluorescence, with red indicating the presence of *D. trenchii*. (**b**) Repeat landscapes shown separately for the CCMP2556 and SCF082 genomes. (**c**) Circle plot depicting the location of syntenic blocks containing collinear gene blocks (i.e. ohnologs) between the CCMP2556 and SCF082 genomes. Ribbons indicate syntenic gene blocks identified with MCScanX that overlap with putative WGD-duplicated regions in both isolates (blue; n=2,427), one isolate only (red; n=612), or neither isolate (black; n=35). (**d**) The percentage of genes in duplicated collinear gene blocks relative to the number of duplicated collinear gene blocks identified within the genomes of Suessiales species. (**e**) Number of genes and syntenic genes recovered for each gene duplication category for the two isolates. (**f**) Phylogenetic tree of Order Suessiales showing the number of lineage-specific gene-family duplications at each node.

### Whole-genome duplication in a coral endosymbiont

We generated *de novo* genome assemblies from *D. trenchii* CCMP2556 (total length = 1.71 Gb; N50 = 774.26 kb) and *D. trenchii* SCF082 (total length = 1.64 Gb; N50 = 398.48 kb) using 10X Genomics linked reads (Tables S1 and S2). The two genomes are highly similar in terms of whole-genome sequence (Table S3, ∼99.7% shared identity), size (Table S4), and repeat landscapes (Fig. 1b, Fig. S1), yielding ∼54,000 protein-coding genes (Table S5) with a comparable level of data completeness to other genome assemblies of Symbiodiniaceae (Table S6; see Methods). To assess WGD in *D. trenchii*, we followed González-Pech *et al.*^11^ to identify collinear gene blocks within each genome (Fig. 1c, see Methods); these blocks likely arose via segmental duplication and/or WGD. We identified 864 blocks implicating 27,597 (49.46% of the total 55,799) genes in CCMP2556, and 776 blocks implicating 18,209 (34.02% of the total 53,519) genes in SCF082 (Tables S7 and S8). The proportion of genes present in collinear blocks in *D. trenchii* is ∼49-fold greater (Fig. 1d) than that in other Symbiodiniaceae and the outgroup dinoflagellate *Polarella*, which have not experienced WGD. We also observed a high extent of conserved synteny (22,041 CCMP2556 genes syntenic with 21,094 SCF082 genes), with ohnologs predominant in these syntenic blocks (CCMP2556: 15,395 [69.85%]; SCF082: 12,617 [59.31%]) (Fig. 1e, Figure S2, and Table S9). Using homologous protein sets derived from available whole-genome data, our inference of lineage-specific duplicated genes (see Methods) revealed 7,945 gene duplication events specific to *D. trenchii*, which is an order of magnitude greater than in other Symbiodiniaceae (Fig. 1f).

Examination of the overall distribution of DNA synonymous substitutions (*K_S_*) showed a distinct peak (Fig. S3), as expected following WGD; the small peak values are explained by the recency of this event in *D. trenchii*^25^. The WGD likely occurred after the split of *D. trenchii* from its sister *Durusdinium glynnii* 0.11–1.93 million years ago (MYA), based on LSU rDNA genetic divergence estimates^5^. Our analysis of whole-genome data following Ladner *et al.*^26^ aligns with these estimates of a Pleistocene origin in the Indo-Pacific (Supplementary Information), a period of frequent sea-level changes in this region^27^. These results, based on independently assembled genomes from two isolates, combined with the extent and size of the gene blocks (Table S7 and Fig. S2), provide unambiguous evidence for WGD in *D. trenchii*.

### Asymmetric divergence of ohnolog-pair expression

To assess putative ohnolog functions in *D. trenchii*, we analysed transcriptome data of CCMP2556^28^ that were generated from free-living cells in culture and from cells in endosymbiosis with the anemone *Exaiptasia pallida*, both under ambient (28°C) and thermal stress (34°C) conditions. We focused on 6,147 expressed ohnolog-pairs that were supported by 10 or more mapped transcripts in ≥50% of the samples, and inferred gene co-expression networks (Fig. S4 and Table S10) using weighted gene co-expression network analysis (WGCNA). Most (4,412 [71.7%] of 6,147) ohnolog-pairs were recovered in different networks, indicating the prevalence of expression divergence between duplicates post-WGD. We then classified ohnolog pairs into five groups based on their differential expression (DE) patterns (Figs 2A, S5-S9; see Methods). Each group exhibited different characteristics (Table S11) relative to expression (Fig. 2a-b), sequence similarity (Fig. S10), gene structure (i.e. exon gain/loss; Fig. 2c), and/or alternative splicing (Fig. S11-S12); see Supplementary Information. Ohnolog-pairs that were differentially expressed between lifestyles observed in only one copy (Group 2; 2,244 [36.5%]%]; Fig. 2d), and those with opposing differential expression observed in any one comparison (Group 5; 100 [1.6%]; Fig. 2e; Table S12) from strongly contrasting expression profiles (most Pearson correlation coefficients < 0; Fig. 2b), showed significantly elevated levels of positive selection, exon gain/loss, sequence divergence, and differential exon usage (DEU; Table S13) relative to the other three groups (all pairwise Wilcox *p* < 0.05; Supplementary Information); these differences were not attributed to the differing number of splice junctions per gene, and ohnologs show greater extent of alternative splicing than singletons (Fig. S13 and Table S14).

**Figure 2.**
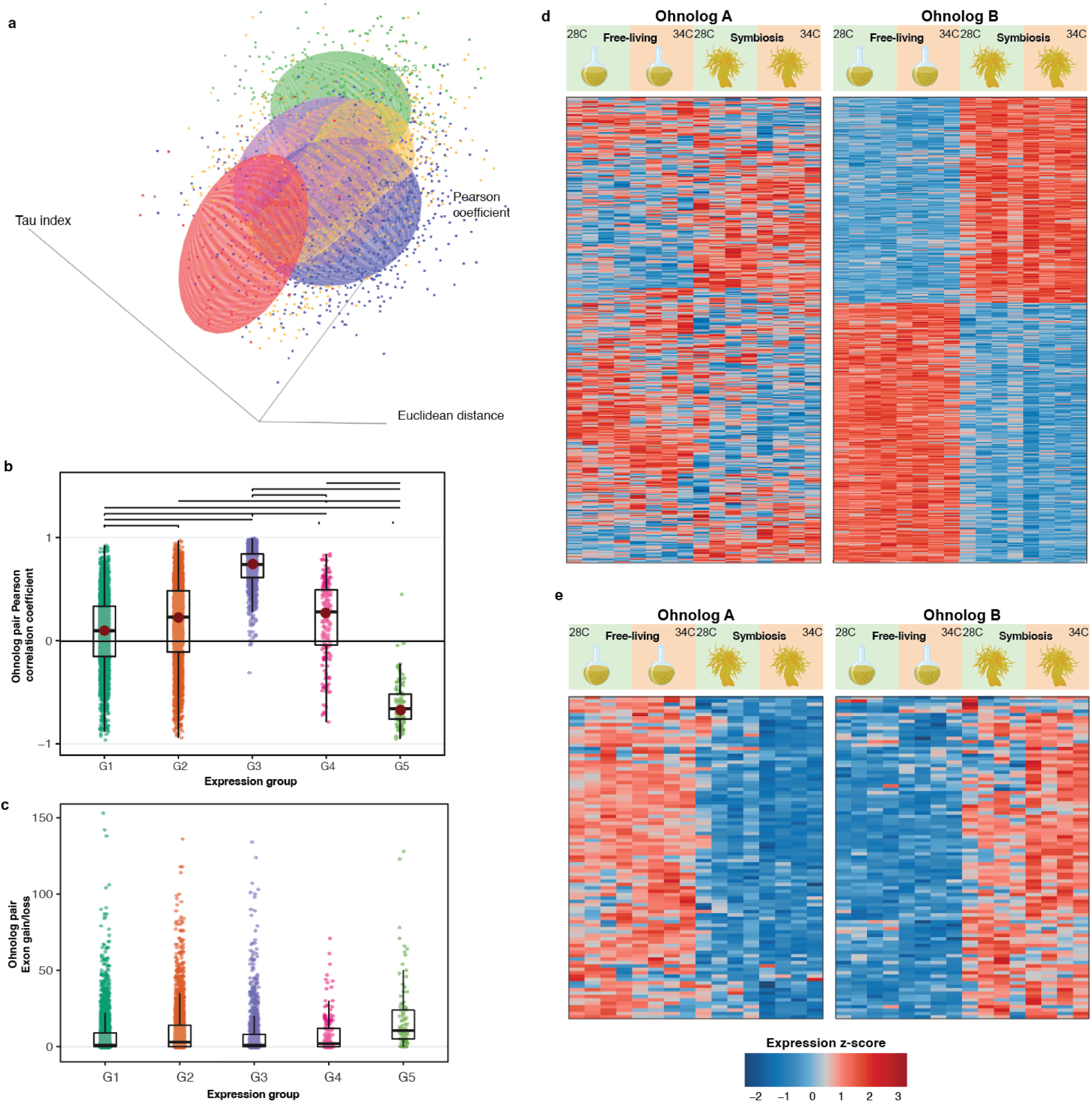
Ohnolog expression post-WGD. (**a**) Three-dimensional scatterplot of the five groups of ohnologs pairs based on their pattern of differential expression (DE), i.e. pairs for which: neither gene showed DE (Group 1; blue); only one showed DE (Group 2; orange); both ohnologs showed DE at the same time in the same manner (Group 3; green); both ohnologs showed DE but at different times (Group 4; purple); and both ohnologs showed DE at the same time but in opposing directions (Group 5; red). (**b**) Pearson correlation coefficients showing correlation of expression patterns between each ohnolog pair within each of the five groups. (**c**) Exon gain/loss between each ohnologs pair within each of the five groups. Heatmaps depicting the normalized gene expression (*z*-score) for (**d**) Group 2 and (**e**) Group 5.

Divergence of expression between ohnologs within a pair can have different outcomes, including the change in expression specificity, an important mechanism for adaptation after WGD. We assessed expression specificity using the τ index^29^ that ranges between 0 (i.e. broad expression, low specificity) to 1 (i.e. narrow expression, high specificity) for all genes that passed the WGCNA quality filtering. We identified 3,508 genes of high expression specificity (τ > 0.7), of which 1,893 (53.96%) were ohnologs (Table S15). Compared to singletons and other duplicate types (except for proximal duplicates), the ohnologs exhibited significantly elevated τ (Fig. S14, Kruskal-Wallis test; *p* < 10^−5^), indicating narrow expression profiles that are more specialized to distinct conditions. This divergence in expression was observed in Group 2 pairs (Fig. S6), for which the differentially expressed copy in each pair showed higher τ and variance in expression relative to its counterpart (Kruskal-Wallis test; *p* < 10^−15^). Whereas most instances of specialized expression are associated with the free-living lifestyle (Table S16), this specialization reflects response to temperatures among the dispersed duplicates (i.e. duplicates separated by >20 genes; Chi-square test post-hoc: *p*=0, residuals=6.15 at 34°C) and the ohnologs (*p* <0.05, residuals=3.10 at 28°C). Interestingly, only ohnologs displayed a tendency towards expression specialization in the symbiotic lifestyle at both 28°C (Chi-square test post-hoc: *p* < 0.01, residuals=3.61) and 34°C (*p* < 0.01, residuals=3.89; Table S16). Although the post-WGD specialization described here relates to lifestyle, parallels are known in multicellular organisms whereby the partitioning of expression across spatiotemporal scales is often observed in different tissues, organs, or developmental stages^14,30^. In post-WGD yeasts, this trend may represent the uncoupling of noise and plasticity in gene expression that enables dynamic gene-expression responses in one of the two duplicates^31^. In *D. trenchii*, this trait may provide greater flexibility in gene expression when cells are free-living or experiencing temperature stress.

The up-regulation or specialization of gene expression by some ohnolog-pairs to different lifestyles in *D. trenchii* appears to either be mediated by, or coincide with, alterations in exon organization. Evolution of WGD-derived genes can lead to loss and/or diversification of alternative spliced forms, and the partitioning of ancestral splice forms between gene duplicates. We investigated this issue by examining the interplay of sequence conservation in exonic sequences with patterns of splice junction conservation and DEU within ohnolog pairs. Based on mean percentage of shared exons per pair, the ohnolog pairs in Group 5 (13.35%) had lower exonic conservation compared to other groups (>16%). DEU across all groups were biased toward functions associated with the free-living lifestyle. Nearly all ohnolog pairs in Group 5 (95 of 100; Fig. 2e and Fig. S9) displayed contrasting differential expression between free-living and symbiotic phases at one or both temperatures, underscoring lifestyle as a strong driver of expression divergence (Supplementary Information). Group 5 ohnologs that are specialized for the symbiotic lifestyle exhibited lower overall DEU, and possessed fewer exons than their counterparts that were up-regulated under the free-living lifestyle (Wilcoxon rank sum test, *p* = 0.015, *V* = 2435.5). These ohnologs also contained exons that were more dominantly expressed during the symbiotic lifestyle (Fig. 3a, Wilcoxon rank sum test, *p* = 0.02798, *V* = 2540). Such a bias in DEU composition towards symbiosis-specialized exons was not observed in the other groups, e.g. Group 2 (Fig. 3b-c). Consequently, the symbiosis-specific DEU, together with the overall decrease in per-gene exons and DEU among symbiosis-associated ohnologs in Group 5, suggest a symbiosis-specific streamlining of gene function. Together with our observation of RNA editing (Tables S17 and S18, Figure S15; Supplementary Information), these results collectively indicate that alterations to gene structure and alternative splicing drive expression divergence of ohnologs in *D. trenchii* that are explained by algal lifestyle.

**Figure 3.**
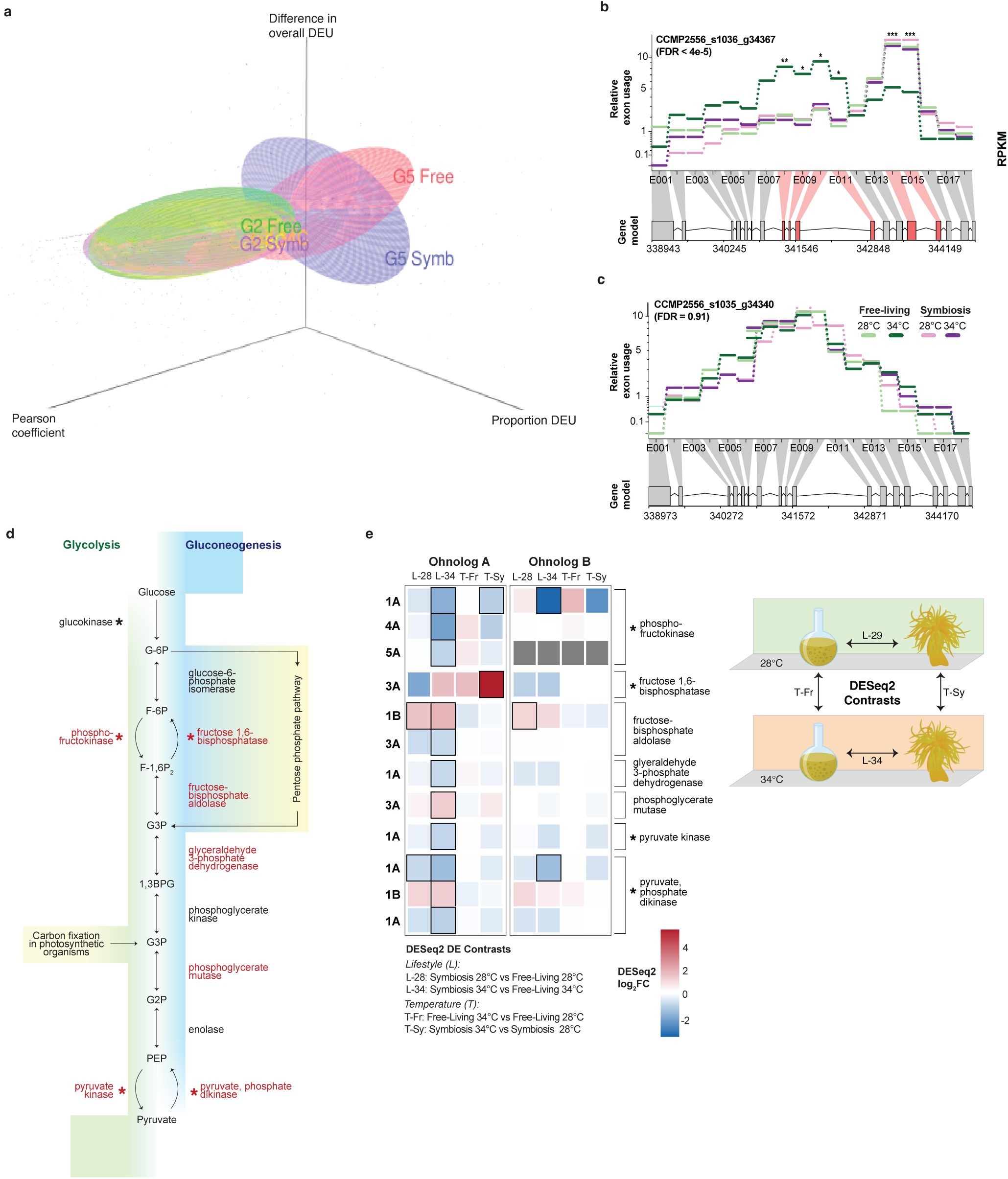
Exon reorganization underlies functional divergence. (**a**) Three-dimensional scatterplot depicting directionality of DEU among ohnologs pairs of Group 2 and Group 5, which reflects the pattern of gene-level DEU. The *z*-axis shows the absolute change in DEU (i.e. overall DEU) within each ohnolog pair from Groups 2 and 5, the *x*-axis shows the relative change in DEU (i.e. proportion of DEU), and the *y*-axis indicates Pearson correlation coefficient of the gene expression. An ohnolog-pair from Group 2 with (**b**) DEU in the ohnolog with gene-level DEU, and (**c**) no DEU in its counterpart. (**d**) Glycolysis and gluconeogenesis pathways for which genes indicated in red were implicated by differentially expressed ohnologs, and an asterisk indicating rate-limiting or key enzymes. (**e**) The log2(fold-change) in expression of differentially expressed ohnologs across distinct comparison of growth conditions with their corresponding scenario indicated on the right.

Although genomic streamlining is usually associated with obligate endosymbionts rather than facultative symbionts, gene duplication may facilitate streamlining in one of the two duplicates in favour of symbiotic lifestyle. Ohnolog pairs of Group 5 were significantly enriched for key functions (Table S19), such as the processing of glutamine and production of the key antioxidant of glutathione, which have been linked to nitrogen cycling associated with symbiosis^32,33^; the implicated genes include glutamine synthetase and S-formylglutathione hydrolase (Table S12). These results suggest that following WGD, specialization of gene expression to distinct conditions may also be enabled by the streamlining of functions and specialization to symbiosis. In contrast, for their duplicated counterparts, the evolution of greater functional flexibility may reflect selection during the free-living phase.

### Partitioned functionality in central metabolic pathways

WGD enables the retention of complete expression networks. Of the 19 inferred co-expression networks (Table S10), different gene duplication types displayed preferential distributions to WGCNA modules (*p* < 2.2 × 10^−16^, *χ*^2^ = 525.63). Singletons and ohnologs were biased towards contrasting co-expression networks, with singletons predominantly associated with networks linked to the symbiotic lifestyle (M1, M8, and M17 in Table S20), and ohnologs with networks linked to a free-living lifestyle (M2, M5, and M6). This result suggests that genes preferentially retained as ohnologs were expressed at contrasting times, compared to those that were lost such that the remaining copies become singletons. Differential expression of ohnologs was observed at the greatest magnitude between lifestyles during heat stress at 34°C (Chi-square test post-hoc: *p* < 10^−3^, Residuals=4.03; Table S21); this may explain in part how *D. trenchii* can establish itself or increase in abundance in new hosts both during and after heat waves^34–38^. These contrasting patterns of singleton and ohnolog membership across co-expression networks indicate a strong association of ohnolog retention with expression networks that are tightly linked to the free-living lifestyle.

We investigated retention of complete metabolic pathways in both *D. trenchii* isolates. Of the 98 pathways retained in duplicate (Table S22), specialization driven by lifestyle was detected in central metabolic pathways (Figs S16-S23), such as glycolysis/gluconeogenesis (Fig. 3d and Fig. S16). Ohnolog specialization in glycolysis/gluconeogenesis reflects the contrasting functions of this pathway during the symbiotic *versus* free-living phases. That is, a high rate of gluconeogenesis, inferred using ohnolog expression data, supplies glucose for translocation to the coral host during symbiosis, whereas a high rate of glycolysis fuels the energetic needs of free-living cells that tolerate more variable environments^3^. Although most enzymes were encoded by Group 2 ohnologs (for which one gene copy was differentially expressed between lifestyles; Fig. 3e), a key rate-limiting enzyme of gluconeogenesis and the Calvin cycle, fructose 1,6-bisphosphatase, was differentially expressed in response to heat stress in symbiosis. Development of minor or partitioned functionality following WGD has been described in duplicate glycolysis pathways^39^. In yeast, these pathways diverged and became semi-independent, with each specialized for low and high glucose levels^39^. In *D. trenchii*, this might allow fine-tuning of carbon metabolism to the contrasting energetic needs of a dual lifestyle.

### Concluding remarks

Our results provide strong evidence that the dual lifestyle has been a key driver of post-WGD genome evolution in the dinoflagellate *D. trenchii*. Our working hypothesis is illustrated in Fig. 4. Under the null hypothesis of a solely free-living lifestyle, we expect post-WGD adaptations to primarily be driven by fluctuating environmental conditions (e.g., nutrient availability). Under the hypothesis of a dual lifestyle that includes symbiosis, adaptations will also strengthen the maintenance of a stable host-symbiont relationship, and efficient nutrient/metabolite exchange within the coral holobiont. Although our results provide stronger support for the free-living phase as the primary driving force behind post-WGD evolution, both lifestyles impact the maintenance and expression divergence of ohnologs. These combined selective forces increase overall fitness in *D. trenchii*, with the greater expression divergence of ohnologs under elevated temperatures a contributor to the high thermotolerance of this species when it is in symbiosis with corals^40^. Benefits conferred by WGD to a free-living lifestyle in more-variable environments, as well as tailoring of post-WGD duplicates to different lifestyles, primed *D. trenchii* to persist longer in the coral holobiont when faced with thermal stress. Whether symbiosis may also have negative effects on fitness post-WGD is unknown^41^. It should be noted that the dual lifestyle is widespread in Symbiodiniaceae^5^, but WGD is not. Although other facultative symbionts within Symbiodiniaceae (e.g., *Cladocopium thermophilum*)^42,43^ are also known for their thermotolerance, WGD was not implicated in these lineages^11,44^. Therefore, the key feature of *D. trenchii* that we are addressing is not dual lifestyle alone, but rather how the capacity for dynamically switching between the symbiotic *versus* free-living phase impacts post-WGD genome evolution and adaptation. Because Symbiodiniaceae propagate to very high densities in coral tissues (10^−5^–10^−6^ cells/cm^2^)^45,46^, the symbiotic phase of *D. trenchii* allows a rapid increase in the population size, particularly of fast-growing genotypes, while resident in host tissues. Consequently, genotypes that have faster growth rates or greater resilience to heat due to WGD-derived adaptations can re-seed free-living populations upon dissociation from the coral due to colony death, bleaching, or other mechanisms of symbiont population control. Repeated cycles of symbiosis followed by the free-living phase may therefore increase the overall fitness of *D. trenchii* populations under the dual lifestyle^47^. Retention of multiple gene copies combined with fixed, adaptive changes likely makes *D. trenchii* more capable of metabolic maintenance under dynamic, often stressful environments, and hence a more-resilient symbiont. Such factors may in turn explain the large geographic and expanded host range of *D. trenchii*^24^ and its well-known capacity for increasing coral survival under heat waves. Therefore, in an intriguing and unexpected twist, WGD, primarily driven by selection under a free-living life phase has converted *D. trenchii* into a coral symbiont able to protect the host coral from thermal stress during symbiosis. *D. trenchii* is also a valuable model for studying the genome-wide impacts of facultative lifestyles.

**Figure 4.**
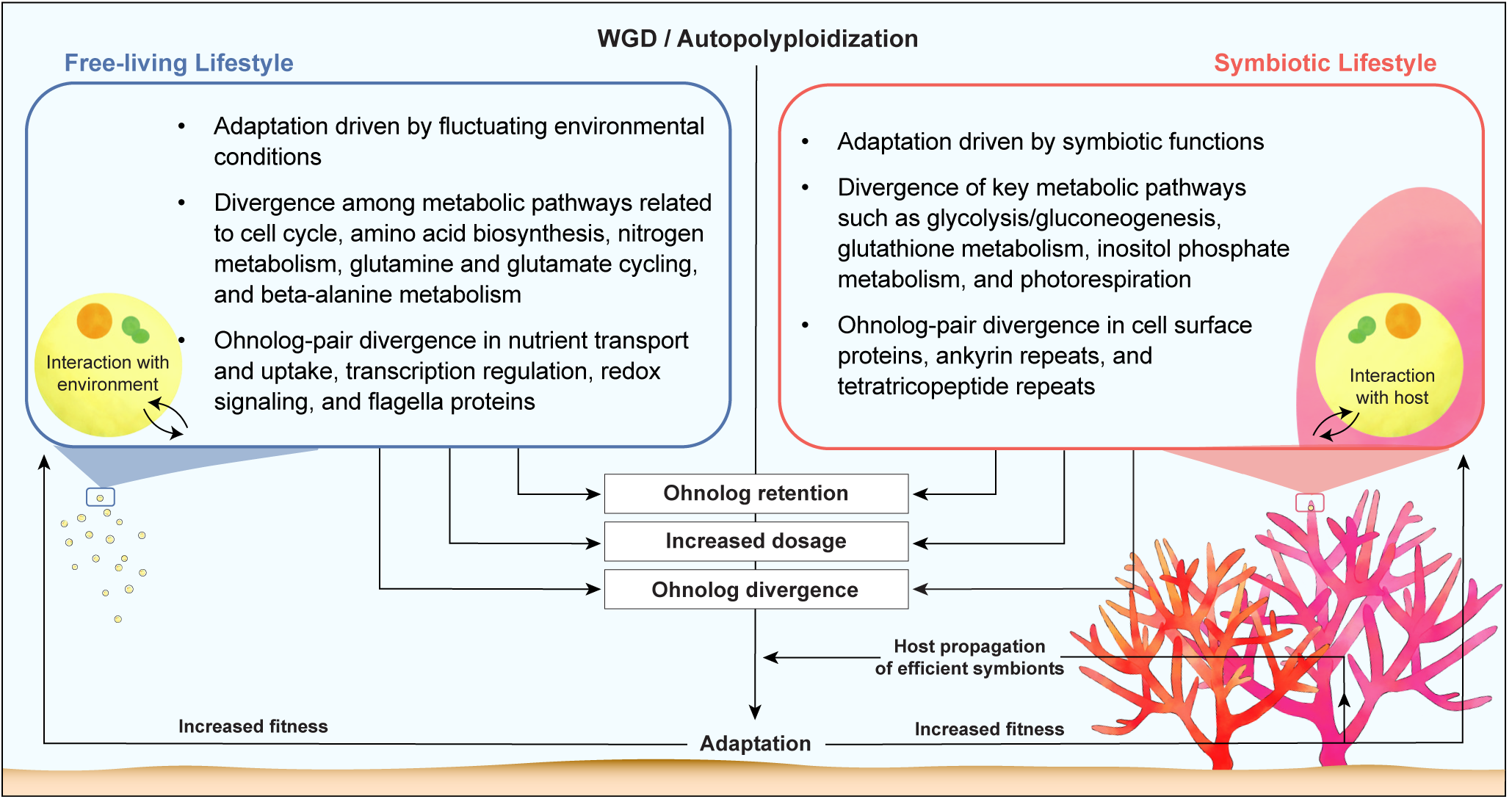
Model of post-WGD divergence in a facultative endosymbiont. Putative selective constraints faced by free-living and symbiotic Symbiodiniaceae under the dual lifestyle are shown, with a focus on post-WGD ohnolog sequence divergence and differential gene expression.

## Methods

### *De novo* genome assembly and prediction of protein-coding genes

*Durusdinium trenchii* strains CCMP2556 and SCF082 (previously designated UTSD amur-D-MI) originally isolated from an *Orbicella faveolate* and *Acropora muricata* coral colonies, respectively, were each separately cultured and genomic DNA extracted for genomic sequencing (see Supplementary Information). Chromium libraries were generated for 10X linked-read sequencing and yielded a total of 236.45 Gbp for CCMP2556 and 212.03 Gbp for SCF082. We assessed the ploidy of *D. trenchii* using *k*-mers and GenomeScope2^48^, which revealed a distinctive single peak in both isolates indicating a haploid genome as seen in other Symbiodiniaceae (Figure S24), with a predicted heterozygosity of 0.31% and 0.20% in CCMP2556 and SCF082, respectively.

For each isolate, a preliminary draft genome was assembled *de novo* using 10X Genomics Supernova v2.1.1. For CCMP2556, the estimated genome coverage (∼100×) exceeded the optimal range (38–56×) of the Supernova assembler; we subsampled the 1.6B reads to 600M reads (∼60× coverage). For SCF082, coverage estimates were observed to be impacted due to the presence of contaminant DNA from microbial sources in the sequencing reads; the *de novo* assembly was generated using all 1.4B reads with the flag *–accept_extreme_coverage*.

Presence of putative contaminant scaffolds in the supernova assemblies was investigated using a comprehensive approach adapted from Iha *et al.*^49^ informed by read coverage, G+C content, taxonomic designation, and *de novo* transcriptome mapping. Taxon-annotated G+C-coverage plots (Figure S25) were generated using the BlobTools suite v1.1^50^ to identify scaffolds in each assembly that deviated by read coverage, taxonomic sequence similarity, and/or G+C content.

Read coverage was assessed using BWA v0.7.17, based on mapping of quality-trimmed reads (Longranger v2.2.2^51^ ran at default setting) to the genome assembly. The taxonomic identity of scaffolds was assigned based on BLASTN search (*E* ≤ 10^−20^) against genome sequences from bacteria, archaea, viruses, and alveolates in the NCBI nt database (release 2021-05-10). *De novo* transcriptome assemblies were mapped to the genome assemblies using minimap2 v2.18^52^ within which we have modified the codes to account for non-canonical splice sites of dinoflagellates. Scaffolds that were designated as non-dinoflagellate were removed from the assemblies if they lacked mapped transcripts from the corresponding *de novo* transcriptome assembly, or when <10% of mapped transcripts indicate evidence of introns in the genomes. We considered a scaffold as a putative contaminant if (a) its sequence coverage or G+C content is not within the 1.5 × interquartile range, and (b) it lacks any transcript support defined above. Upon removal of these putative contaminant sequences from the CCMP2556 assembly, the filtered assembly was incorporated in the database as the *D. trenchii* reference for assessing the assembled scaffolds of SCF082 using the same approach.

Publicly available RNA-Seq data from previous studies of CCMP2556^28^ and SCF082^53^ were used to further scaffold the assembled genome sequences (see Supplementary Information). RNA-Seq reads for both isolates were first quality-trimmed using fastp^54^ (mean Phred quality ≥ 30 across a 4bp window; minimum read length of 50bp). For each isolate, the filtered reads were assembled *de novo* using Trinity v2.11.0^55^ independently for each treatment. The transcriptome assemblies for CCMP2556 (791,219 total transcripts) and those for SCF082 (355,411 total transcripts) were mapped to the filtered genome assemblies using minimap2 v2.18^52^ that was modified to recognize the non-canonical splice sites of dinoflagellates. The mapped transcripts were then used to scaffold the filtered genome assemblies with L_RNA_Scaffolder^56^ at default parameters.

A second round of scaffolding was then performed with ARBitR^57^, which incorporates the distance information from linked-read sequencing data when merging and scaffolding assemblies. Longranger BASIC quality-trimmed linked genome reads (outputs from the standard 10X Genomics data workflow) were mapped to the scaffolded genome assemblies for ARBitR scaffolding, yielding the final genome assemblies: CCMP2556 (assembly size = 1.70 Gb; N50 = 750 Kb; 29,137 scaffolds) and SCF082 (assembly size = 1.64 Gb; N50 = 398.5 Kb; 44,682 scaffolds) (Table S2). The CCMP2556 assembly is the most contiguous reported in Symbiodiniaceae aside from the recent chromosome-level assemblies for *Symbiodinium microadriaticum*^58^ and *Breviolum minutum*^59^.

Genome and gene features of dinoflagellates are highly idiosyncratic and atypical of eukaryotes, due in part to non-canonical splice sites^60^. Therefore, the prediction of protein-coding genes from dinoflagellate genomes requires a comprehensive workflow (https://github.com/TimothyStephens/Dinoflagellate_Annotation_Workflow/) tailored for these features, guided by high-confidence evidence^61^. Here, we adopted a customised workflow integrating the results from multiple methods, guided by available transcript and protein sequences, independently for CCMP2556 and SCF082; see Supplementary Information for detail.

### Analysis of whole-genome duplication

We first searched for evidence of collinear gene blocks using MCScanX^62^ in intra-species mode (*-b 1*) to identify putative duplicate gene blocks within each genome (i.e. segmental and/or whole-genome duplication), and in inter-species mode (*-b 2*) to identify syntenic gene blocks between the two genomes. A collinear block is defined as at least five genes conserved in the same orientation and order as a result of segmental duplication and/or WGD events. For each comparison, all-vs-all BLASTP search results were restricted to the top five hits (query or subject coverage > 50%; *E* ≤ 10^−5^). Predicted genes from each genome were classified using *duplicate_gene_classifier* (within MCScanX) into singleton, dispersed duplicates (i.e. duplicates separated by >20 genes), proximal duplicates (i.e. duplicates separated by <20 genes), tandem duplicates, and WGD/segmental duplicates (i.e. ohnologs).

Second, we assessed the reconciliation between each gene tree and the species tree; the topological incongruence between the two trees indicates history of gene duplication or loss^63^ OrthoFinder v2.3.10^−64^ was first used to infer homologous gene sets among Suessiales species using BLASTP (*E*-value ≤ 10^−5^). Multiple sequence alignments were performed with MAFFT v7.487^65^ (*-linsi*), from which phylogenetic trees were inferred using FastTree v2.1.11^66^ at default parameters. Reconciliation of gene-tree and species-tree within OrthoFinder was then used to identify lineage-specific duplication events; those specific to *D. trenchii* indicative of WGD-derived duplicated genes (i.e. ohnologs).

Third, we assessed the impact of WGD on the rate of synonymous substitution (*K_s_*) among all homologous gene sets, using CCMP2556 as the reference, following the wgd pipeline^67^. Briefly, homologous protein clusters were inferred using a Markov Clustering algorithm^68^ from the previous all-versus-all BLASTP search (used for MCScanX), and aligned using MAFFT^65^. Phylogenetic tree for each homologous protein cluster was inferred using FastTree2^66^ and used to estimate *K_s_* values for each cluster using codeml implemented in PAML^69^. A Gaussian-mixture model was applied to the *K_s_* distribution, using a four-component model that provided the best fit for the data according to Akaike information criterion (AIC), yielding a final node-averaged histogram of *K_s_* distribution. To estimate the timing of WGD, we first calculated the estimated substitution rate (*r*) per year in Symbiodiniaceae adapting the approach of Ladner *et al.*^26^ to incorporate genome data and the updated divergence time estimates from LaJeunesse *et al.*^5^.

We followed Aury *et al.*^20^ to infer metabolic pathways that were preferentially retained in duplicate following WGD using PRIAM v2 (January 2018 release). Briefly, we identified metabolic enzymes that had been uniquely retained as ohnologs or singletons. We then compared the proportion of enzymes uniquely retained as ohnologs to singletons, to the background proportion of the number of ohnologs and singletons annotated in the genome. This tests whether the number of uniquely retained metabolic enzymes for a particular pathway exceeds the background levels that would be expected to occur by random. We additionally required (a) five or more distinct enzymatic proteins to be identified as uniquely retained in either duplicate or singleton, and (b) pathways to be significantly overrepresented in both isolates. The proportion of enzymes coded by genes that were uniquely retained as ohnologs or singletons, compared to their overall proportions in the genome, was used to determine which KEGG pathways were preferentially retained in duplicate following WGD.

### Evolution of ohnolog expression

Trimmed RNA-Seq reads (above) were mapped to the corresponding genome using HISAT2 v2.2.1 (*--concordant-only*) with a Hierarchical Graph FM index informed by annotated exon and splice sites. Counts of uniquely mapped paired-end (PE) reads overlapping with CDS regions were then enumerated using *featureCounts* (*-p --countReadPairs --B -C*) implemented in Subread v0.2.3^70^. The raw counts were filtered to remove lowly expressed genes using the *filtrByExpr* function in edgeR. Differential gene expression analysis was performed with edgeR using a generalized linear model. We considered genes to be differentially expressed when false discovery rate (FDR) < 0.01 and the absolute value of log_2_(fold-change) > 1. We compared the difference between lifestyles at two temperatures, i.e. symbiosis *versus* free-living at 34°C (L-34), and symbiosis *versus* free-living at 28°C (L-28), and the response to temperature stress in the two lifestyles, i.e. 34°C *versus* 28°C at free-living (T-Fr), and 34°C *versus* 28°C in symbiosis (T-Sy).

A weighted gene co-expression network analysis (WGCNA) was performed on all genes in R using the WGCNA package. Variance of normalised counts were calculated using the standard DESeq2 workflow followed by its *varianceStabilizingTransformation*. Because symbiosis is a strong driver of expression in Symbiodiniaceae, using the inferred soft-thresholding power for reducing noise and setting a required threshold for gene correlations would have yielded a mean connectivity of over 4,000 at the inferred power of 6. Therefore, a weighted, unidirectional co-expression network was inferred using a power of 18, the recommended value for signed networks with less than 20 samples. Co-expression modules were inferred using the function *blockwiseModules* that collectively infers signed networks (networkType=“signed”, TOMtype=“signed”, maxBlockSize=10000, corType=“bicor”, maxPoutliers=0.05, pearsonFallback=“individual”, deepSplit=2, dcuth=0.999, minModuleSize=30, reassignThreshold=0.1, cutHeight=0.2).

We calculated the adjacencies using a signed network with *bicor* robust correlation coefficient (power=18, type=“signed”, corFnc=bicor, maxPoutliers=0.1, pearsonFallback=“individual”). A topological association matrix was then inferred with a signed network and *dissTOM* computed from the product. A hierarchical dendrogram of genes was inferred using *hclust* (method=“average”). The dendrogram was cut using *cutreeDynamic* (deepSplit=2, minClusterSize=15, cutHeight=0.999) and the cut dendrogram merged with *mergeCloseModules* (cutHeight=0.15, corFnc=bicor, maxPoutliers=0.1, pearsonFallback=“individual”). Preferential distribution of the different gene duplication categories to WGCNA modules was assessed with a chi-square test and a post hoc analysis performed with the R package *chisq.posthoc.test* (https://github.com/ebbertd/chisq.posthoc.test.git).

Expression specificity of ohnologs was assessed using the tau (τ) index^29^, where τ = 1 indicates highly specific expression, and τ = 0 indicates broad expression. The log-normalised fragments per kilobase million (FPKM) counts were used to calculate τ index scores for those genes with a log_2_(FPKM + 1) > 1 in at least one condition following Yanai *et al.*^29^. The τ indices for the different MCScanX duplication categories were compared using a Kruskal-Wallis rank sum test; pairwise comparisons using Wilcoxon rank sum test with continuity correction and holm *p*-value adjustment were performed to determine differences between the duplication categories. A chi-square test of all significant τ indices (τ ≥ 0.7) was conducted to assess potential biases in expression specificity for treatments among the duplication categories.

### Analysis of post-transcriptional regulation

All-versus-all BLASTN search (query or subject coverage > 50%; *E* ≤ 10^−20^) was used to identify shared exonic sequences that have been retained since WGD. For inferring differential exon usage (DEU) within genes among the treatment conditions, gene models were first broken up into exon “counting bins” using the Python script *dexseq_prepare_annotation.py* from DEXSeq. The relative usage of each exon bin, i.e. the number of transcripts mapping to the bin or to the gene, was then calculated from the HISAT2 BAM file using *dexseq_count.py*. The DEXSeq R package was then used to infer differential exon usage within genes using a generalised linear model, correcting for significance at the gene level using the Benjamini-Hochberg method^71^.

To examine the conservation of splice junctions in ohnolog pairs, all *de novo* assembled transcripts were first aligned to the genome using a minimap2 v2.20^52^ with code modified to recognize alternative splice sites in dinoflagellates, from which splices sites were identified and annotated using PASA^72^. Splice sites categorized as alternative acceptor, alternative donor, alternative exon, retained exon, and skipped exon were retained for subsequent analysis. Each identified splice event was assigned two unique identifiers to represent the upstream and downstream positions of the splice event, along with its gene identifier and genomic location. The upstream and downstream 300bp-region for each splice event were then extracted using the bedtools v2.30 *flank* and *getfasta* functions. An all-*versus*-all BLASTN search of the extracted splice junction sequences was used to identify sequence similarity (*E* ≤ 10^−5^) between the sequences. Custom Python scripts were used to filter the BLASTN results to identify conserved splice junctions, in which both upstream and downstream regions for a splice event in two ohnologs were significantly similar (*E* ≤ 10^−5^). The splice junction profile for each ohnolog pair was then converted to a binary representation, where the presence of a splice junction in an ohnolog was represented as 1 and the absence of a splice junction represented as 0 (i.e., conserved splice junctions represented as 1 in both ohnologs compared to 0 for those that were not conserved). A Kendall’s rank correlation was then conducted in R to identify ohnolog pairs that exhibited high level of conservation in splice junctions. An exact binomial test was also performed to identify ohnolog pairs that had diverged in terms of total splice junctions (*p* < 0.05).

## Data availability

The genome data generated from this study for the two *D. trenchii* isolates are available at NCBI through BioProject accession PRJEB66001. The assembled genomes, predicted gene models, and proteins for *D. trenchii* CCMP2556 and SCF082 are available at https://doi.org/10.48610/27da3e7.

## Acknowledgements

This Project was supported by funding from the Australian Research Council grants DP190102474 awarded to C.X.C. and D.B. and FL180100036 to M.J.H.v.O., the Australian Academy of Science Thomas Davies Grant for Marine, Soil, and Plant Biology awarded to C.X.C., the National Science Foundation grant CAREER1453519 awarded to M.R.L., the City University of New York grant PSC-CUNY 69757-0047 awarded to C.G.C., the National Institute of Food and Agriculture-US Department of Agriculture Hatch grant NJ01180 awarded to D.B., and funding from the University of Technology Sydney to T.K. and D.J.S. This project is also supported by computational resources of the National Computational Infrastructure (NCI) National Facility systems through the NCI Merit Allocation Scheme (Project d85) awarded to C.X.C. We thank Brian Kemish and David Green for their technical assistance during genome assembly as well as Joel Burke and Michelle Havlik for technical assistance with the algal cultures.

## Author contributions

K.E.D., A.J.B., D.J.S., C.X.C., and M.R.L. conceptualized the study; K.E.D., D.B., and C.X.C. drafted the initial manuscript with A.J.B., M.J.H.v.O., D.J.S., and M.R.L. helped finalize the text; A.J.B. and R.M.A. maintained the cultures and extracted genomic DNA samples; K.E.D. performed all bioinformatic analyses except for the analyses on RNA editing and organellar genomes done by Y.C. and S.S., respectively. Y.C. performed RNA editing analysis and S.S. organellar genome analysis. All authors read and approved the manuscript.

