## Supplementary Information for "Whole-genome duplication in an algal symbiont bolsters coral heat tolerance"

### 1 Introduction to *Durusdinium trenchii*

#### 1.1 Symbiodiniaceae and coral reef health

The health of coral reef ecosystem is critically regulated by the functional performance of the dinoflagellate microalgae (Family Symbiodiniaceae) that thrive in endosymbiosis with reef-forming corals. Optimum metabolically derived nutrient exchange between these microalgae and coral[1,2] fuels coral growth and calcification that are essential to reef productivity and accretion. However, anthropogenic stressors are increasingly driving sub-optimum environmental conditions for symbiosis that result in an accumulation of toxic metabolic byproducts (e.g. Reactive Oxygen Species[3]), and in turn endosymbiont rejection from the host ("coral bleaching"). Episodes of mass, region-wide coral bleaching have become increasingly common place, changing the entire nature of coral reef systems[4,5]. Importantly, Symbiodiniaceae taxa are extremely diverse taxonomically[6] and functionally[7]. Taxa that are more stress tolerant, or rather, those suited to more suboptimal reef environment conditions, can proliferate in coral tissues and enable persistence under conditions where corals would otherwise bleach[8,9], including present day "natural extremes", such as super-hot tidal pools and lagoons[10,11]. Resolving how these stress tolerant symbionts have evolved and their phenotypes remain a major goal in efforts to better manage reef systems into the future, including through more-aggressive active interventions to evolve more-thermotolerant symbionts[12-14] or manipulating host-symbiont partnerships[15,16].

#### 1.2 *Durusdinium trenchii*: the thermotolerant symbiont

Among the Symbiodiniaceae taxa, the genus *Durusdinium* includes coral symbionts known to survive in large variations of temperature and turbidity in nature[6]. Earlier evidence for whole-genome duplication (WGD) in *Durusdinium trenchii* based on microsatellite studies[17-19] was linked to its thermotolerance[20]. The notion of WGD driving the evolution of enhanced thermal tolerance is widely supported by eco-physiological studies showing *D. trenchii* often replaces the predominant coral symbiont taxa following thermal stress, thereby conferring increased thermal tolerance to the host[21-23]. In general, greater thermotolerance of *D. trenchii,* the relatively recent spread of this species in the Caribbean[19] demonstrate the potential for *D. trenchii* to enhance resilience of corals to warming oceans.

#### 1.3 Evolutionary significance of whole-genome duplication

Polyploidy can promote the success of mutualistic associations[24,25]. For instance, higher colonisation rates of the mycorrhizal fungi in the roots has been linked to polyploidy in some plants[25]. Conversely, the link between higher stress tolerance and ease of invasion by polyploids has been connected to a greater likelihood of being an invasive species[26,27], and can have cascading effects on community interactions and structure[28].

Duplicated genes derived from a WGD event are commonly referred to as the *ohnologs*, and are believed to provide the raw genetic material for evolutionary innovation[29,30]. WGD promotes the specialisation of ohnolog expression or the partitioning of expression across spatiotemporal scales, often discussed in context of different developmental stages[31], tissue types[31-33], or under biotic/abiotic stress conditions. The biological implications of diverging ohnolog expression has been examined extensively in whole-genome studies of polyploid organisms with WGD including plants[34,35], amphibians[32], and fishes[33].

The preservation of ohnologs after WGD in a genome can be manifested in three ways: increased dosage, neo-functionalisation, and sub-functionalisation[36,37]. Increased dosage relates to the increased number of genes encoding the same, likely critical function. In neo-functionalisation, the ohnologs evolved to gain new functions, potentially as an adaptive mechanism to increase fitness[38]. In sub-functionalisation, the ohnologs evolved to encode different parts of their ancestral function. This is thought to occur via two models: the escape from adaptive conflict (EAC) model[39], and the duplication, degeneration, complementation (DDC) model[40]. The EAC model represents scenarios where each sub-function that is ultimately preserved cannot be simultaneously optimised, requiring adaptive changes to fix different sub-functions of the ancestral function to escape the conflict. On the other hand, the DDC model pertains to scenarios where successive neutral mutations remove different subsets of the ancestral function over time in a complementary fashion, resulting in diverged ohnologs with complementary functions. The processes that underpin these two models are not mutually exclusive, and the specific mechanism for retention of ohnologs can be dynamic over time[41].

### 2 Generation and analysis of genome data

#### 2.1 Culture maintenance of *D. trenchii*

The *Durusdinium trenchii* strain CCMP2556 was originally isolated by Dr Mary Alice Coffroth (Buffalo University, New York, USA) from an *Orbicella faveolata* coral sampled at 10m depth on Tennessee Reef, Florida Keys (24.7335° N, 20.7669° W). The CCMP2556 culture was first acquired from the National Center for Marine Algae and Microbiota (Bigelow Laboratory for Ocean Sciences, East Boothbay, Maine), and maintained for one year in PROV-50 medium (NCMA) with an antimicrobial cocktail (kanamycin [50 µg/mL], ampicillin [100 µg/mL], streptomycin [50 µg/mL], amphotericin B [2.5 µg/mL]) to minimise bacterial and fungal contamination. During this time, the culture was passaged in the antimicrobial cocktail and inspected via light microscopy to confirm culture quality and the absence of microbial contamination on a monthly basis. PCR amplification and sequencing of the *cp23S* marker sequence was also regularly conducted for comparison to the known *D. trenchii* sequence to verify the genetic identity of the cultures. No other *Durusdinium* spp. were maintained in the laboratory prior to or during this time. To prepare the CCMP2556 cells for extraction of genomic DNA, bacterial contamination was minimized following methods adapted from Su *et al.*[42]. Briefly, $5\times10^7$ log-phase cells were pelleted via centrifugation (500 *g*, 3 min) and resuspended in sterile PROV-50 medium (40 mL) using the antimicrobial cocktail above. Cells were then pelleted and resuspended in PROV-50 medium with 0.005% Tween-80 and EDTA (0.1 M, pH 8.0). Resuspended cells were incubated (25°C, 1 h) on a platform rocker followed by the addition of lysozyme (0.5 mg/mL) and SDS (0.25%) and another incubation with rocking at room temperature (RT; 10 min). Finally, cells were washed three times with PROV-50 medium containing the antimicrobial cocktail, and snap-frozen in liquid nitrogen prior to DNA extraction.

The *D. trenchii* strain SCF082 (previously designated UTSD amur-D-MI) was originally isolated from an *Acropora muricata* coral colony on the Great Barrier Reef (Magnetic Island) and maintained at the Australian Institute for Marine Science (AIMS) culture collection, prior to shipment to University of Technology Sydney. SCF082 was grown in xenic cultures in 200mL IMK medium (Daigo's IMK powder for microalgae) for 2 weeks at 24ºC under 25 µmol/s/$m^2$ light and pelleted in 50mL Falcon tubes (ELMI CM-6MT centrifuge ~184 *g*). To reduce bacterial contamination, cell pellets were washed three times by re-suspending them in sterile PBS buffer (50 mL) and pelleted (ELMI CM-6MT centrifuge ~184 *g*). Bacterial contamination was assessed before and after wash based on plating on Marine Agar, and cell count by flow cytometry: cells were stained with SYBR® Green (Merck) in the dark (15 min) prior to detection by a CytoFLEX S flow cytometer (Beckman Coulter) at 488 nm and bandpass filter 525/40. Flow cytometer results showed a 90% reduction in bacterial contamination. Final pellets were combined and pelleted at 4000 *g* (5 min), and snap-frozen in liquid nitrogen before DNA extraction.

#### 2.2 DNA extraction

To extract genomic DNA from CCMP2556 cells, frozen cells (above) were ground together with acid-washed 425-600 µm glass beads (Sigma) in a liquid nitrogen-chilled mortar and pestle. The ground cells (~50 mg) were used for DNA extraction using the MagAttract HMW DNA Kit (Qiagen) following the manufacturer's protocol. To minimise DNA degradation, wide-bore pipette tips were used for all steps involving the handling of lysate or DNA, and mixing was performed by gentle inversion instead of the manufacture-indicated pipetting. The final DNA was eluted in Buffer AE (Qiagen; 10 mM Tris-Cl 0.5 mM EDTA, pH 9.0) and stored at -20°C. The DNA sample was analysed using pulse-field gel electrophoresis which indicated the presence of an intact sample (mean fragment size ~100 kb), confirming DNA integrity. The 10X Chromium library was constructed using 1 ng of genomic DNA as input following the manufacturer's protocol.

To extract genomic DNA from SCF082, cell lysis was performed on snap-frozen cells using a custom-made French press with two passes at 6895 kPa. Lysate was collected and subjected to standard extractions using ice-cold phenol:chloroform:isoamyl alcohol (25:24:1, v/v/v), and chloroform:isoamyl alchohol (24:1, v/v). DNA was precipitated using sodium acetate (pH 5.2; final concentration 0.3 M), and an equal volume of ice-cold isopropanol. Precipitated DNA was washed with 70% ethanol and resuspended in TE buffer (RT, overnight) to allow for high-molecular weight fragments to redissolve. DNA was size selected using PluePippin (SageScience) pulse-field gel electrophoresis prior to 10X Chromium library preparation following the manufacturer's protocol.

#### 2.3 Genome sequencing and assembly

##### 2.3.1 Linked-read genome sequencing

The *D. trenchii* CCMP2556 Chromium library was sequenced at HudsonAlpha Institute for Biotechnology (Huntsville, AL) using the Illumina HiSeq X platform (two lanes, paired-end 2×150 bases). The *D. trenchii* SCF082 Chromium library was sent to Ramaciotti Center for Genomics, University of New South Wales (Sydney, Australia) for sequencing using Illumina NovaSeq 6000 (S1 flow cell, paired-end 2×150 bases). Genome sequencing yielded a total of 236.45 Gbp for CCMP2556 and 212.03 Gbp for SCF082 (Table S1).

##### 2.3.2 *De novo* genome assembly and contaminant removal

See the Methods section in main text for details of *de novo* genome assembly and the removal of putative contaminant sequences.

##### 2.3.3 Genome scaffolding using transcriptome data

Publicly available RNA-Seq data from previous studies of CCMP2556[43] and SCF082[44] were used to further scaffold the assembled genome sequences. The CCMP2556 RNA-Seq data[43] were generated at ambient temperature and under heat stress, each for the two lifestyles: free-living in culture and in symbiosis with the anemone *Exaiptasia pallida.* Free-living cells and symbiotic anemones were maintained at 28°C, with the temperature of heat stress treatments ramped up 2°C per day for 3 days to a final temperature of 34°C. Four biological replicates were then sampled from each of the four treatments (free-living at 28°C, free-living at 34°C, symbiosis at 28°C, and symbiosis at 28°C) on day 3. The SCF082 RNA-Seq data[44] were generated from free-living cells in culture at two temperature levels, both ambient temperature and heat stress, and at two timepoints. Free-living cells were maintained at 26°C, with heat stress treatments ramped up 2°C per day for 3 days to a final temperature of 32.4°C, and then maintained at the temperature for seven days. Four biological replicates were sampled from control and heat stress treatments on day 3 after initial temperature ramping and at day 10 after 7 days of heat stress.

RNA-Seq reads for both isolates were first quality-trimmed using fastp[45] (mean Phred quality ≥ 30 across a 4bp window; minimum read length of 50bp). For each isolate, the filtered reads were assembled *de novo* using Trinity v2.11.0[46] independently for each treatment. The transcriptome assemblies for CCMP2556 (791,219 total transcripts) and those for SCF082 (355,411 total transcripts) were mapped to the filtered genome assemblies using minimap2 v2.18[47] that was modified to recognize the non-canonical splice sites of dinoflagellates. The mapped transcripts were then used to scaffold the filtered genome assemblies with L_RNA_Scaffolder[48] at default parameters. A second round of scaffolding was then performed with ARBitR[49]. See the Methods section in main text for details of the linked-read scaffolding.

#### 2.4 Genome size and ploidy

We first assessed the ploidy of *D. trenchii* using *k*-mers and GenomeScope2[50], which revealed a distinctive single peak in both isolates indicating a haploid genome (Figure S24). Genome size was estimated following the approach by Lin *et al.*[51], and inferred using *k*-mers enumerated from filtered genome-sequence reads. Briefly, the quality-trimmed reads were mapped to the final genome assembly using HISAT2 v2.2.1[52] (*--very-sensitive*). The *k*-mers were then extracted from these mapped reads using Jellyfish v2.3.0[53] at $k$ = 19, 21, 23, 25, 27, 29 and 31, from which a genome size estimate was calculated independently for each *k*. The average value of these seven estimates yielded the final genome size estimate: 1.05 Gb for CCMP2556 and 1.65 Gb for SCF082 (Table S4); these sizes may be underestimated due to high extent of repetitive or duplicated genomic regions.

#### 2.5 Identity confirmation and sequence comparison of genome data

The identity of the two isolates was confirmed by the presence of identical ribosomal large subunit sequences in both genomes, and near-identical ITS2 sequences (i.e. one mismatch in the 639bp-region of D1/D2) between them. Sequence similarity between the CCMP2556 and SCF082 genomes was further assessed using the nucmer function (--mum) implemented in MUMmer v4.0.0[54], requiring alignments of 100bp, 1Kb, and 10Kb with a minimum sequence identity at 90%.

### 3 Genome Annotation

#### 3.1 Repetitive elements

Repetitive sequences were first masked in the genomes prior to gene model prediction. A *de novo* repeat library was generated in RepeatModeler v2.0.2 (http://www.repeatmasker.org/RepeatModeler/) and combined with known repeats (RepeatMasker database release 20181026) for repeat masking in RepeatMasker v4.1.2 (http://www.repeatmasker.org/). We then used the RepeatMasker Perl scripts *calcDivergenceFromAlign.pl* to calculate Kimura distances for each repeat type and *createRepeatLandscape.pl* to graph the genome repeat landscapes.

Compared to other Symbiodiniaceae taxa[55], the repeat landscapes of both *D. trenchii* genomes (Figure 1b and Figure S1) show a distinct distribution of predominantly LTR repeats, of which many are highly conserved (those constituting ~2% of the genome sequences have Kimura distances centered between 0 and 5). Long-interspersed nuclear elements (LINEs) and some LTRs appear to be highly divergent (Kimura distances >25), lending support to the notion of ancient LINE expansion in Symbiodiniaceae and *Polarella*, pre-dating the diversification of Suessiales[55,56].

#### 3.2 Customised workflow for *ab initio* prediction of protein-coding genes

Genome and gene features of dinoflagellates are highly idiosyncratic and atypical of eukaryotes, due in part to non-canonical splice sites[57]. Therefore, the prediction of protein-coding genes from dinoflagellate genomes requires a comprehensive workflow (https://github.com/TimothyStephens/Dinoflagellate_Annotation_Workflow/) tailored for these features, guided by high-confidence evidence[58]. We adopted a customised workflow integrating the results from multiple methods, guided by available transcript and protein sequences, independently for CCMP2556 and SCF082.

##### 3.2.1 Gene prediction using transcriptome and protein evidence

Genome-guided transcriptome assemblies were first generated to complement the previous *de novo* transcriptome assemblies (**Section 2.3.3**). To do this, quality-trimmed RNA-Seq data were mapped to the genome assembly using HISAT2 v2.2.1 (*--no-discordant*) and assembled with Trinity v2.11.0 under the genome-guided mode (*--genome_guided_max_intron 70000*). For each of the four growth conditions, the *de novo* and genome-guided transcriptome assemblies (total of 8 assemblies across all conditions) were first "cleaned" with SeqClean (https://sourceforge.net/projects/seqclean/) and the UniVec database. The transcripts were mapped to the genome with BLAT[59] (*-maxIntron=70000*) and the resulting alignments validated for accuracy in PASA v2.4.1[60] modified to recognize dinoflagellate non-canonical splice donor sites (*--MIN_PERCENT_ALIGNED=75, --MIN_AVG_PER_ID=95, --NUM_BP_PERFECT_SPLICE_BOUNDARY=0*). Transcripts that did not pass validation were then re-mapped using a minimap2 (for which the code was modified to account for alternative splice sites)[57], and the minimap2 alignments validated using the same parameters as the BLAT analysis. Valid BLAT and minimap2 alignments were then used as inputs (*--IMPORT_CUSTOM_ALIGNMENTS_GFF3*) to generate an initial set of predicted genes using the customized PASA v2.4.1 (for which the code was modified to account for alternative splice sites), and coding sequences (CDSs) predicted with TransDecoder v5.5.0[61]. Available 689,426 predicted protein sequences from Suessiales taxa and all protein sequences from Swiss-Prot release 02032020 were combined with the custom repeat library used in RepeatMasker to generate a second set of gene predictions using MAKER v3.01.03 in the *protein2genome* mode[62].

##### 3.2.2 *Ab initio* gene prediction

We performed *ab initio* gene prediction using three methods: the unsupervised method using GeneMark-ES[63], and the supervised methods of AUGUSTUS[64] and SNAP[65]. Genome sequences with soft-masked repeats (from Section 3.1) were used for gene prediction in GeneMark-ES v4.65[63]. For AUGUSTUS and SNAP, we used the transcript-based gene model predictions from PASA (**Section 3.2.1**) to first generate a high-quality training set. Briefly, we identified gene models with near full-length CDSs using BLASTP search (query coverage ≥ 80%; $E \leq 10^{-20}$) to a combined database of RefSeq non-redundant proteins (release 98) and available Suessiales proteins (134,488,139 proteins total). We identified and removed any sequences that share similarity to transposable elements in the JAMg transposon database (https://sourceforge.net/projects/jamg/files/databases/) using HHblits v3.3.0 (probability = 80%; $E$-value = $10^{-5}$) and TransposonPSI (http://transposonpsi.sourceforge.net/), and further reduced sequence redundancy using cd-hit v4.8.1[66] (*-c 0.75 -n 5*). The final set of high-quality gene models (i.e. the "golden genes") was then identified from the remaining sequences using the Perl script *prepare_golden_genes_for_predictors.pl* within the JAMg pipeline (http://jamg.sourceforge.net/), modified to recognize GA donor splice sites. These "golden genes" were used as a training set in *ab initio* gene prediction with SNAP[65] and AUGUSTUS v3.4.0[67], both were customized to recognize the non-canonical splice sites.

##### 3.2.3 Integration of multiple gene predictors

We used EvidenceModeler v1.1.1[60], modified to account for non-canonical splice sites in dinoflagellates, to integrate the results from the multiple methods (i.e. PASA, MAKER, AUGUSTUS, SNAP, and GeneMark-ES) into a comprehensive set of evidence-based predictions. The weightings used for the different gene sets (PASA 10, MAKER protein 8, AUGUSTUS 6, SNAP 2 and GeneMark-ES 2) leveraged transcriptome- and protein-based sources more heavily. We quality-filtered the integrated EvidenceModeler predictions to only keep gene predictions that were supported by at least two *ab initio* prediction programs, or have transcript-based support (i.e. predicted by PASA).

#### 3.3 Gene model refinement and functional annotation

The resulting gene models (**Section 3.2.3**) were further refined using the validated PASA alignments from the initial gene prediction (**Section 3.2.1**) to correct for exon-intron boundaries, incorporate alternative splice forms, and identify 5′- and 3′-untranslated regions. Four rounds of gene-model refinement were performed, with the refined predictions from each round used as the input for the subsequent round of refinement. This yielded our final set of gene models for both *D. trenchii* isolates, with 55,799 protein-coding genes for CCMP2556, and 53,519 for SCF082. Gene models were functionally annotated based on BLASTP ($E \leq 10^{-5}$) searches against the UniProtKB (release 2021_03; Swiss-Prot and TrEMBL) database. Pfam domains within predicted proteins were annotated using PfamScan ($E \leq 0.001$) in conjunction with the Pfam-A database.

Completeness of the genomes and the predicted proteins was assessed against the Benchmarking Universal Single-Copy Ortholog (BUSCO) Alveolata (alveolata_odb10) dataset with BUSCO5 v5.1.2[68]. The higher percent recovery of complete BUSCOs in the predicted proteins of *D. trenchii* CCMP2556 (74.8%) and SCF082 (82.4%) than in most published dinoflagellate genomes indicates the high quality and completeness of the predicted gene models (Table S6). The recovery of complete BUSCOs was also higher than the 52.1% completeness of the earlier published draft genome of *D. trenchii* from isolate NIES-2907[69], which also represents the smallest Symbiodiniaceae genome assembly, with the fewest predicted genes. We excluded the NIES-2907 assembly from further analyses due to these differences, and the fact that the NIES-2097 assembly, generated using only short-read sequencing data, would not adequately resolve genomic regions implicated by WGD.

### 4 A shared whole-genome duplication in *D. trenchii*

We used three methods to examine putatively WGD-derived genes and confirm WGD in *D. trenchii*: (1) identification of within-genome and between-genome collinear gene blocks, (2) gene tree-species tree reconciliation, and (3) the distribution of the rate of synonymous substitution ($K_s$). We also estimated the timing of WGD.

#### 4.1 Identification of WGD-derived genes

**Materials and Methods**

We adopted three approaches to identify WGD-derived genes, detailed in the Methods section of the main text. To assess positive selection, the rates of non-synonymous (Ka) and of synonymous (Ks) substitutions for each duplicate gene pairs was calculated using the add_ka_to_ks_to_synteny.pl (https://github.com/reubwn/microhomology) Perl script, based on the pairwise alignment generated using Clustal Omega v1.2.474.

To estimate the timing of WGD, we first calculated the estimated substitution rate (*r*) per year in Symbiodiniaceae adapting the approach of Ladner *et al.*[70] to incorporate genome data and the updated divergence time estimates from LaJeunesse *et al.*[6]. Orthologous protein families were inferred using OrthoFinder v2.3.10[71] and BLASTP (*-evalue 1e-5*) using representative proteins from the genomes of six genera. For each family, the protein sequences were aligned using MAFFT v7.487 (*linsi* mode) and trimmed using trimAl[72] by specifying back-translation (*-automated1 -ignorestopcodon -backtrans*) to obtain the trimmed codon alignment. Estimates for $K_s$ were then obtained using custom Perl scripts with CODEML from PAML. The maximum tree length cutoff (number of substitutions per codon) was conservatively set at 12 or twice the number of taxa. The $K_s$ estimates from trees passing this cutoff were then used to estimate the substitution rate ($r$) and calculate the estimated timing of WGD, $T = \frac{K_s}{2r}$, where $K_s$ represents the average rate of synonymous substitutions in WGD-derived gene blocks (i.e. collinear gene blocks).

#### Results and Discussion

We identified a substantial level of conserved synteny between the two genomes, with 3,070 syntenic blocks implicating 22,041 (39.50% of the total 55,799) genes in CCMP2556 and 21,094 (39.41% of the total 53,519) genes in SCF082 (Figure 1d). The syntenic regions span the majority of both genomes yet in the more contiguous regions of the genome (i.e. longer scaffolds), suggesting a lack of power in more-fragmented regions due to lower quality. Our identification of within-genome collinear gene blocks indicated substantial support for WGD (Tables S7 and S8) based on 864 blocks implicating 27,597 (49.46% of the total 55,799) genes in CCMP2556, and 776 blocks implicating 18,209 (34.02% of the total 53,519) genes in SCF082 (Figure 1e). Importantly, the synteny of these putative ohnologs appears highly conserved between the two genomes, i.e. 55.8% of 27,597 CCMP2556 ohnologs versus 69.3% of 18,209 SCF082 ohnologs (Table S9), and not limited to a particular genomic region (Figure 1c). There was sufficient sequence divergence for calculating $K_a/K_s$ values in 7,797 of the 13,811 ohnolog pairs within CCMP2556, and 5,011 of the 9,272 pairs within SCF082. Although these values indicate a similar level of positive selection in the two genomes (i.e. percentage of ohnolog pairs with $K_a/K_s > 1$ is 68.50% for CCMP2556 and 68.63% for SCF082), they also indicate divergent selective pressures between syntenic ohnolog pairs recovered in both genomes ($p < 2e\text{-}16$, $R^2 = 0.08313$).

Considering microsatellite studies implicated a potential segmental or small-scale duplication in the *D. trenchii* and *D. glynnii* lineage prior to WGD in *D. trenchii*[73], we assessed whether any support for this small-scale duplication event was present within the duplicated collinear gene blocks. Focusing within the more-contiguous genome assembly of CCMP2556, we identified only 20 genes in more than one collinear block due to the assembly contiguity in contrast to SCF082 where many were found in multiple collinear blocks. Of these 20 genes in CCMP2556, 18 are involved in a potential tetrad as they were implicated in a single collinear block independently in four scaffolds (CCMP2556_s118, CCMP2556_s119, CCMP2556_s2202, CCMP2556_s2203), whereas the corresponding syntenic collinear block in SCF082 was only recovered in two scaffolds (SCF082_s839 and SCF082_s840). Although this observation does not exclude the potential for a small-scale duplication in the lineage leading to *D. trenchii* and *D. glynnii* prior to WGD in *D. trenchii*, further investigation with *D. glynnii* is needed to confirm this. The number and size of the collinear gene blocks along with their near-perfect representation in duplicates promotes a scenario of WGD as the source of the duplication. It is unlikely that this observation was caused by many, non-overlapping segmental duplications.

Our analysis of gene tree-species tree reconciliation revealed an order of magnitude greater gene-family duplication in *D. trenchii* than the other genera of Symbiodiniaceae (Figure 1f). We identified 7,945 protein families with *D. trenchii*-specific duplication events, implicating 17,241 (62.5%) and 16,025 (88.0%) ohnologs respectively in CCMP2556 and SCF082. The equivalent number of protein families is <1000 for other genera, e.g. 554 for *Symbiodinium* and 959 for *Cladocopium* (Figure 1f). results supports the notion of WGD in the evolution of *D. trenchii*, distinct from other Symbiodiniaceae taxa.

Synonymous substitutions ($K_s$) accumulate in genes at specific rates according to the organism, and can be employed to estimate the timing of WGD as all ohnologs originated from the same WGD will have accumulated synonymous mutations at a similar rate for the same amount of time compared to the rest of the genome. This can be seen as a distinct peak in a distribution and which we to assess for WGD. While we find a distinctive peak only at a relatively small $K_s$ (0.026 for CCMP2556, 0.024 for SCF085) that would indicate WGD in *D. trenchii* (Figure S3).

The small peak of $K_s$ values can be explained by the recent WGD expected in *D. trenchii*[74], which likely coincides with its split from the sister *Durusdinium glynnii* ~1 million years ago[6]. Our independent analysis placed generated a revised substitution rate of $\sim 1.41 - 1.42 \times 10^{-7}$ for the family Symbiodiniaceae following the established approach in Ladner *et al.*[70] with genomic data using the revised Symbiodiniaceae divergence estimates from LaJeunesse *et al.*[6], and placed the timing of the *D. trenchii* WGD event between 595.2 and 617.9 Ka (thousand years ago).

While $K_s$ plots are informative for not only inferring but also identifying the age of WGD events, they are also prone to misestimation in both ancient and young WGD events[74,75]. In young WGD events, as seen here, the relatively limited time may not have been sufficient for the two duplicate genes to diverge in sequence similarity. This makes the discrimination between WGD duplicates and small-scale duplicates near impossible, as both distributions of synonymous substitution would represent that of exponential decay. However, our results based on whole-genome data from two *D. trenchii* isolates, and the extensive duplication of syntenic collinear gene blocks supported by large-scale gene-family duplications revealed by gene tree-species tree reconciliation analysis, provides strong support for WGD in the evolutionary history of *D.*

*trenchii*. Interestingly, the majority of syntenic ohnolog-pairs displayed similar $K_a/K_s$ ratios from their respective genomes (3,570 [65.98%]), with twice as many (1,897 [43.08%]) under positive selection than purifying selection (22.89%). However, $K_a/K_s$ ratios suggest a third of syntenic ohnolog-pairs are under contrasting selective regimes with 1,416 having $K_a/K_s$ ratios > 1 in CCMP2556 only, and 1,403 with $K_a/K_s$ ratios > 1 in SCF082 only.

Dinoflagellates typically have a haploid, motile vegetative phase (which we studied), therefore, the *k*-mer spectrum (Fig. S24) we report is expected for the *D. trenchii* strains. The strong evidence for massive gene families and many large segmentally duplicated regions shared by the two strains, however, indicates that a WGD (i.e., a polyploidisation event) occurred in the common ancestor of these lineages. High sequence identity among genome segments within and between strains suggests autotetraploidy. We postulate that selection for genome reduction, likely due to the dual lifestyle, led to subsequent massive genome reduction that resulted in extant haploid genomes with the comparative data illuminating the polyploid history. These cells therefore provide models for post-WGD gene and genome evolution that increase fitness under the dual lifestyle.

### 5 Facultative lifestyle drives ohnolog-pair expression specialization

As WGD-derived genes (i.e. ohnologs) can exhibit both conservation and partitioning of functions and expression, we examined the relative contributions and drivers of each to post-WGD adaptation in *D. trenchii*. We assessed this in context of the facultative lifestyle and temperature, both for which *D. trenchii* is known through its vast geographic and expanded host range as well as notorious thermotolerance. We evaluated expression through three approaches: (1) similarity in expression profiles, (2) contribution of ohnologs to differential responses of gene expression, and (3) level and direction of expression specificity. We then compared expression to measures of sequence similarity, exon gain/loss, and positive selection to determine what aspects of gene structure are linked to expression divergence.

#### 5.1 Global expression divergence

**Materials and Methods**

A primary mechanism for the emergence of evolutionary novelties following WGD is the duplication of entire signaling networks, including metabolic pathways. The duplication of these entire pathways and networks enables them to subsequently diverge in concert. Following the formula used in Aury *et al.*[76], we inferred metabolic pathways that were preferentially retained in duplicate following WGD; see Methods for detail.

We examined how the expression of signaling networks might have diverged post-WGD by inferring global co-expression networks, and weighted gene co-expression network analysis (WGCNA) to assess co-expression of genes; this analysis was performed on all genes in R using the WGCNA package; see Methods for detail.

Expression specificity of *D. trenchii* ohnologs was assessed using the tau (τ) index[77], where $\tau = 1$ indicates highly specific expression, and $\tau = 0$ indicates broad expression. The log-normalised fragments per kilobase million (FPKM) counts were used to calculate τ index scores for those genes with a $\log_2(\text{FPKM} + 1) > 1$ in at least one condition following Yanai *et al.*[77]. The τ indices for the different MCScanX duplication categories were compared using a Kruskal-Wallis rank sum test; pairwise comparisons using Wilcoxon rank sum test with continuity correction and holm *p*-value adjustment were performed to determine differences between the duplication categories. A chi-square test of all significant τ indices ($\tau \geq 0.7$) was conducted to assess potential biases in expression specificity for particular treatments among the duplication categories.

$$\text{Tau index: } \tau = \frac{\sum_{i=1}^{n}\left(1 - \widehat{X}_\iota\right)}{n-1};\ \widehat{X}_\iota = \frac{x_i}{max_{1\leq i\leq n}(x_i)}$$

We then assessed the functional role of ohnolog-pairs within the broader transcriptional response of *D. trenchii* to changes in temperature and lifestyle. We used edgeR to identify differentially expressed (DE) genes (FDR < 0.05, $\log_2$[fold-change] > 0.5), and examine the relative contributions of ohnologs to their overall differential response compared to genes classified in other duplication categories. We then grouped ohnolog pairs according to the co-occurrence, directionality, and extent of gene expression change.

#### Results and Discussion

WGD enables the duplication of entire signaling networks, the function of which can subsequently diverge in concert in relation to environmental conditions. This is particularly true for autopolyploids that emerge as a result of hybridization. We assessed metabolic pathways in

*D. trenchii* that were retained in duplicate, and if they exhibit divergence in expression of the implicated genes. In total, we identified 132 metabolic pathways that were preferentially retained in duplicate in both isolates (Table S22), including central pathways such as glycolysis/gluconeogenesis (Figure 3d-e), cell cycle, spliceosome, and ubiquitin mediated proteolysis.

We then assessed for the extent of similarity, or any biases, in co-expression network membership present in ohnologs in gene expression co-expression networks as a proxy for large-scale functional divergence in expression. One common approach for this is the inference of co-expression clusters or modules using algorithms[78-80] or specific programs such as WGCNA[81], and then examining the distribution of duplicate gene pairs, or ohnologs pairs, throughout the clusters or modules. To this end, we employed stringent filtering prior to WGCNA analysis, 35,052 genes were ultimately included the WGCNA analysis including 5,857 singletons, 10,996 dispersed duplicates, 294 proximal duplicates, 759 tandem duplicates, and 17,146 ohnologs.

Of the 19 inferred co-expression networks (Table S10), different gene duplication categories displayed preferential distributions among the WGCNA modules (Chi-square test: $p < 2.2e\text{-}16$, $\chi^2 = 525.63$). Examination of the residuals from the chi-square post-hoc test (Table S20) showed that the different categories of gene duplication displayed variable membership preferences. The three co-expression modules (i.e. M1, M8 and M17) in which singletons were more prevalent were associated with symbiotic lifestyle, with M1 and M8 linked to elevated temperatures and M17 to ambient temperature. Ohnologs were more prevalent in co-expression modules of M2, M5 and M6 that are associated with a free-living lifestyle, with the strongest M6 linked to elevated temperatures during a free-living lifestyle and the remaining two with ambient temperatures. Further, two modules linked to singletons (M1, M8) and the three linked to ohnologs (M2, M5 and M6) were negatively associated with each other. These contrasting patterns of singleton and ohnolog membership across co-expression networks suggests that genes preferentially retained as ohnologs and those returned to singleton status are involved in different signaling networks, with ohnolog retention associated with expression networks more tightly linked to a free-living lifestyle.

Compared to singletons and ohnologs, all other duplicate categories displayed a lesser extent of preferential module membership. Both tandem and dispsersed duplicates were negatively associated with co-expression networks connected to symbiosis (M1), whereas dispersed duplicates, similar to ohnologs, were positively associated with those related to a free-living lifestyle (M3). Proximal duplicates displayed preferential membership in M16 that was associated with both free-living lifestyle and temperature stress.

This bias towards belong to co-expression networks linked to greater activity during a free-living lifestyle was further reflected within the differential gene expression analysis (Figs. 2d-e), where ohnologs were uniquely biased towards being differentially expressed between the two lifestyles during heat stress (Chi-square test: $p < 1e\text{-}3$, Residuals=4.03, Table S21), in contrast to the other gene types particularly dispersed duplicates which were significantly less likely to be DE (Chi-square test: $p < 3e\text{-}3$, Residuals=-3.80). Further, ohnologs displayed a strong bias against DE in response to heat stress both when free-living (Chi-square test: $p < 7e\text{-}3$, Residuals=-4.14) and in symbiosis (Chi-square test: $p < 4e\text{-}3$, Residuals=-3.75). This further supports a scenario where ohnolog expression has become specialized to a particular lifestyle and changes its expression to respond to the changing environment during lifestyle transitions.

We identified 3,508 genes with high expression specificity ($\tau \geq 0.7$), of which 1,893 (53.96%) were ohnologs. Ohnologs had a significantly elevated $\tau$ (Kruskal-Wallis test; $p < 1e\text{-}5$) compared to singletons and other duplicates except for the 462 proximal duplicates, with which there was no significant difference. This suggests ohnologs tended to be more specialized in expression to distinct conditions and have lower expressional breadth than most genes. We further examined the timing of these narrowed expression profiles by looking at when the expression was maximal, revealing a dominance in the free-living lifestyle (i.e. Free-living 24°C: 624 ohnologs and Free-living 34°C: 684 ohnologs, compared to Symbiosis 28°C: 228 ohnologs and Symbiosis 34°C: 357 ohnologs), and to a lesser extent during symbiotic heat stress.

Altogether, these results support a scenario where ohnolog retention post-WGD was preferentially retained in metabolic pathways or signaling networks that are more highly used during a free-living lifestyle. Considering the high expression specificity in the free-living lifestyle and during symbiotic heat stress response, along with the direction of regulation of these DE ohnologs, our results suggest that the specialization of ohnolog expression was primarily toward a free-living lifestyle, although this remains to be validated.

#### 5.2 Patterns of ohnolog-pair co-expression

**Materials and Methods**

To identify potential links between patterns of positive selection and the divergence of ohnolog sequences and expression, we assessed pairwise sequence similarity of ohnologs in each Clustal Omega alignment (**Section 4.1**), the associated exon gain/loss (i.e. absolute difference in exon number), and the $K_a/K_s$ estimates (**Section 4.1**) relative to ohnolog-pair expression divergence (**Section 4.1**). Overall expression divergence was assessed by calculating the Euclidean distances of their $log_2$FPKM expression and assessed with a Mann-Whitney test of $log_2$FPKM. Statistical significance was assessed through pairwise comparisons using Wilcoxon rank sum tests with continuity correction. Kendall correlation coefficients were calculated from the expression of the two ohnologs in a pair and used in a linear regression to assess the connection between pairwise sequence similarity or exon gain/loss (difference in total number of exons) to overall expression distance. Outliers were identified using Cook's distance[82]; those with a distance at least three-fold greater than the mean were removed. The relative importance of exon gain/loss and sequence similarity was assessed using the Lindeman, Merenda and Gold (LMG) method for variance decomposition[83] in the *calc.relimp* function from the *relaimpo* R package[84]. Expression variance between the two ohnologs in a pair was assessed using F tests and of $log_2$FPKM expression values with the function *var.test* from the *stats* package in R.

We then assessed the functional role of ohnolog-pairs within the broader transcriptional response of *D. trenchii* to changes in temperature and lifestyle. We used edgeR to identify DE genes (FDR < 0.05, log2FC > 0.5), and examine the relative contributions of ohnologs to the overall DE transcriptional response compared to other gene duplication categories. We examined the direction and co-occurrence of DE ohnolog pair to better understand the specific roles performed by ohnolog-pairs that have experienced distinct changes in expression profiles. The ohnolog pairs were then grouped according to the co-occurrence, directionality, and extent of gene expression change.

**Results and Discussion**

Expression divergence of duplicate gene pairs was first proposed by Ohno[29] as the first step in the functional divergence of duplicate genes, also increasing their likelihood of retention and fixation in the genome. We first examined this in regard to the WGCNA module membership of ohnolog pairs, in which most paired ohnologs were recovered in different co-expression modules, suggesting a greater extent of divergence than conservation of expression between them. There were 6,147 ohnolog-pairs (1,737 in same module; 4,410 in different modules) classified into 19 co-expression modules. Those duplicate pairs that belong to the same modules are presumably less likely to have diverged in function and more likely to have retained their ancestral functions compared to those in different modules, and might relate to the specialization to lifestyle.

We then investigated the features that potentially drive or coincide with expression divergence of ohnologs. These features include the accumulation of mutations (i.e. sequence divergence) and changes to gene structure in terms of exon gain/loss (i.e. absolute difference in exon number); the latter may be a stronger driver of expression divergence and specialization in recent WGD[85]. This showed that overall sequence similarity and exon gain/loss explained 18.8% of the variance in expression; of this variance, 27.6% was attributed to sequence similarity, and 72.4% to exon gain/loss. This result aligns with that of other young WGD events such as in goldfish, where changes to gene structure and specifically the gain/loss of exons is believed to have enabled more rapid functional divergence than changes to sequence similarity. Although sequence similarity of ohnolog-pairs was greater in those belonging to the same co-expression module (Kruskal-Wallis test; $p = 0.013$; $W = 6.19$), exon gain/loss (Kruskal-Wallis test; $p < 1e{-}11$; $W = 46.77$) and $K_a/K_s$ levels (Kruskal-Wallis test; $p < 1e{-}12$; $W = 40.34$) were both stronger predictors of whether an ohnolog pair are in the same co-expression module. Previous studies using transcriptome data have documented a higher level of positive selection in the *D. trenchii* lineage compared to other Symbiodiniaceae genera[70]; this may be driven in part by functionally divergence of WGD-derived ohnologs that are experiencing elevated levels of positive selection.

Differential expression analysis of ohnolog-pairs revealed five main patterns in the DE responses of ohnolog pairs (Figure 2a). In order of prominence, these groups represent ohnolog pairs where neither displayed any DE (Group 1; 2,515 pairs), only one displayed DE (Group 2; 2,244 [36.5%] pairs), both showed similar patterns of DE (Group 3; X pairs), both were DE but in response to different conditions (Group 4; X pairs), and both displayed contrasting DE to the same condition (Group 5; 100 [1.6%] pairs). Most ohnolog-pairs (4,147 [71.7%]) belonged to different co-expression networks, suggesting divergence of expression within ohnolog-pairs was a likely scenario since WGD; only ohnolog pairs in Group 3 are more likely to share similar co-expression pattern. The contrasting patterns of co-expression was primarily driven by ohnolog pairs for which both copies were not differentially expressed (Group 1), those where only one was DE (Group 2; Figure 2d), and those that were DE in contrasting patterns in response to the same condition (Group 5; Figure 2e). Patterns of sequence similarity, exon gain/loss, exon usage, and selective pressure varied among the groups (Table S11), with Groups 2 and 5 displaying significantly greater rates of exon gain/loss (Figure 2b) and positive selection ($K_a/K_s$) compared to the others.

Between each of the 2,236 ohnolog-pairs in Group 2, only one copy was DE. This pattern may represent a collection of scenarios. A potential explanation may relate to trade-offs between noise and plasticity after small- and large-scale gene duplication events such as WGD, as previously described in yeasts[39]. A potential explanation may relate to trade-offs between noise and plasticity after small- and large-scale gene duplication events such as WGD. For instance, in yeasts[39], genes that function in both stress response and cellular maintenance may be unable to simultaneously optimise its functionality that is specific to stress response. Gene duplication relieves these effects, and enables one of the two duplicates to display dynamic expression and respond rapidly to a changing environments, while the other displays a more-stable expression profile. The DE ohnolog from these pairs displayed not only more-narrow expression profiles, but also higher variance and higher τ than the non-DE ohnolog (Kruskal-Wallis test; $p < 1e\text{-}15$; $H= 560.96$). For the DE ohnologs in these pairs, their greater expression specificity to the free-living lifestyle at both temperatures, and to symbiotic lifestyle under heat stress, hints at the potential directionality of this specializiation. Another potential explanation might be that the non-DE ohnolog is in the process of pseudogenisation.

Interestingly, for each of the 100 ohnolog pairs in Group 5, both copies were DE in the same condition but in opposing directions. This pattern appears to be strongly dependent on the factor of lifestyle: 13 pairs between lifestyles under ambient temperature, 61 pairs between lifestyles at elevated temperature, and 21 pairs between lifestyles at both temperatures. In contrast, this pattern was observed in only one pair between ambient and elevated temperatures in the free-living lifestyle, and in the remaining four pairs between temperatures in the symbiotic lifestyle. Combined with the highest $K_a/K_s$ ratio (mean=3.31, median=1.55), the greatest exon gain/loss ratio (mean=19.0, median=10.5), and the lowest pairwise sequence identity (mean=93.1%, median=98.1%) relative to all groups, these 100 pairs in Group 5 present clear evidence for strong functional divergence and specialization to different lifestyles.

### 6 Post-transcriptional regulation

#### 6.1 RNA editing

**Materials and Methods**

The editing of mRNAs in CCMP2556 was inferred using JACUSA2 with HISAT2-mapped RNA-Seq reads and bowtie2-mapped longranger-trimmed 10X genomic reads (above) using the *--very-sensitive* mode. PCR duplicates were removed from the BAM files using the *MarkDuplicates* function from the Picard Toolkit v2.26.2 (https://broadinstitute.github.io/picard/). Putative sites of mRNA editing identified by JACUSA2 were first filtered to meet the criteria of the D, Y, H feature filters. A second round of filtering was then performed using the R package JACUSA2helper, further requiring a score ≥ 1.15, >10 read coverage in DNA, >5 read coverage in each RNA replicate, observed bases ≤ 2, and the inference of RNA editing in all four treatments. We then examined ohnolog divergence by searching for evidence of differential mRNA editing in relation to symbiotic lifestyle (i.e. edited site exclusive to either lifestyle) and/or temperature stress (i.e. edited site exclusive to 28°C or to 34°C).

**Results and Discussion**

Editing of mRNAs is a post-transcriptional regulation that can diversify an organism's response beyond what is coded in the genome. It has been described in dinoflagellate genomes[86], but is yet to be examined in context of WGD. In the CCMP2556 genome, we identified 3,754 mRNA edited sites in gene regions: 1,802 implicated in synonymous substitutions, 1,909 in missense (non-synonymous) mutations, and 43 in nonsense mutations (e.g. premature stop codons or truncation of protein sequences). Of these, most edited sites were identified in dispersed duplicates (1,351 sites), followed by ohnologs (1,255 sites), singletons (500 sites), tandem duplicates (92 sites), and proximal duplicates (33 sites; Table S17). Although 1,138 ohnolog pairs exhibit RNA editing in one of the two ohnologs, very few (199) pairs showed RNA editing in both ohnologs.

Among genes expressed in cells grown in the four distinct conditions, we identified 3,754 (free-living at 28°C), 2,643 (symbiosis at 28°C), 2,687 (free-living at 34°C), and 2,654 (symbiosis at 34°C) mRNA-edited sites. Of the total sites, more than one-half (1,112 [54.8%]) were identified in all four treatments, but all the edited sites were identified in genes expressed free-living at 28°C (Figure S15); none were exclusively found in symbiotic or free-living cells grown at 34°C. These results suggest that the majority of mRNA editing in *D. trenchii* occurs when living freely outside a host, or in symbiosis in the absence of temperature stress.

*D. trenchii* CCMP2556 displayed minimal overall differential RNA editing when compared to other dinoflagellates. Of the 16 genes identified with differential mRNA editing (Table S18), 15 (93.8%) were associated with the symbiotic lifestyle, including a light-harvesting complex stress-related protein 3.1 (LHCSR3.1), sporozoite surface antigen MB2, two peridinin-chlorophyll *a*-binding proteins, and an insulin-degrading enzyme. These symbiosis-driven edited sites were predominantly located in CDS regions; in comparison, the edited sites of two genes, hypoxia-inducible factor prolyl hydroxylase 2 (HIF-PH2) and G-protein-signaling modulator 1 (Gpsm1), were restricted to intron regions. The latter may represent editing in non-coding RNA and/or regulatory regions within introns. Differential mRNA editing between 28°C and 34°C was only found in the intron regions of two genes, Gpsm1 and a reticulocyte-binding protein 2 homolog a.

#### 6.2 Alternative splicing and exon usage

**Materials and Methods**

See the main text Methods for details regarding the analysis of splice junctions conservation in ohnolog pairs.

Common exons within ohnolog pairs were assessed using an all-vs-all BLASTN search (query or subject coverage > 50%; $E \leq 10^{-20}$) to identify shared exonic sequences that have been retained since WGD. For inferring differential exon usage (DEU) within genes among the treatment conditions, gene models were first broken up into exon "counting bins" using the Python script *dexseq_prepare_annotation.py* from DEXSeq. The relative usage of each exon bin, i.e. the number of transcripts mapping to the bin or to the gene, was then calculated from the HISAT2 BAM file using *dexseq_count.py*. The DEXSeq R package was then used to infer differential exon usage within genes using a generalised linear model, correcting for significance at the gene level using the Benjamini-Hochberg method[87].

**Results and Discussion**

One mechanism that can contribute to the diversification and expansion of the proteome post-WGD is alternative splicing. In this context, ohnolog evolution is captured by three models: an independent model where there is no change in the level of alternative splicing, a function-sharing model where the alternative splicing events are partitioned between the ohnologs, and an accelerated model. We assessed this in *D. trenchii* by identifying putative locations of alternative splicing (Table S14), and how they varied among different gene duplication categories. On average, ohnologs contained significantly more (800) splice junctions per gene compared to any other category (each <400; Figure S13). We then compared the alternative splicing profiles within ohnolog pairs to determine if alternative splicing might be implicated in the divergence of gene duplicates following WGD. For each ohnolog pair, we determined the number of intron/exon splice junctions, then assessed the conservation of each splice junction by comparing upstream and downstream regions. We found that although some differed in the total number of splice sites and their within-pair conservation (569 pairs), most differed in either conservation of splice sites (1587 pairs) or the overall number of splice sites (1786 pairs; Table S23). However, we observed no significant pattern among the five expression groups or in relation to the level of positive selection.

We assessed the variability in exon usage due to alternative splicing by combining gene isoforms into non-overlapping exon "counting bins" and compared the number of mapped transcripts per exon across the gene expression treatments. We refer to this index as differential exon usage (DEU). We compared a reduced model that controlled for the other factors as a covariate (i.e. a reduced model for temperature includes lifestyle as a covariate) to identify significant DEU across treatments (Benjamini-Hochberg adjusted $p < 0.025$). We inferred DEU at the gene level (FDR < 0.025) by accounting for the multiple tests of DEU for each exon in a gene. We found significant DEU in 10,857 genes: most (10,341) are in relation to lifestyle, followed by temperature (1,444), with only 7 in both lifestyle and temperature (Table S13). Taken together, the increased number of splice junctions and DEU in ohnologs implicate the divergence of alternative splicing in the evolution of ohnologs following WGD.
