## Supplementary Figures S1-S25 for "Whole-genome duplication in an algal symbiont bolsters coral heat tolerance"

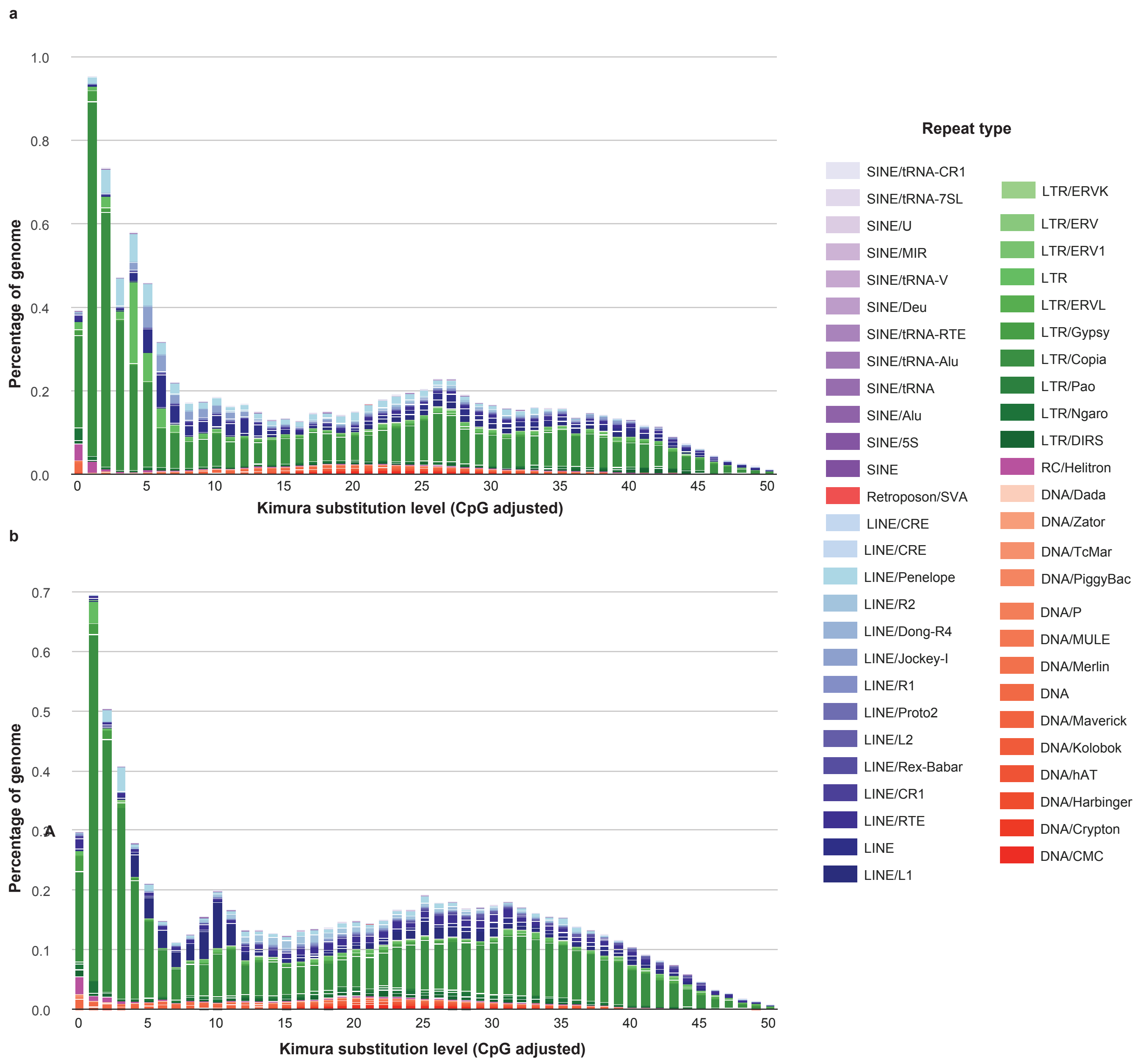


**Figure S1 |** *Durusdinium trenchii* repeat landscape distributions.

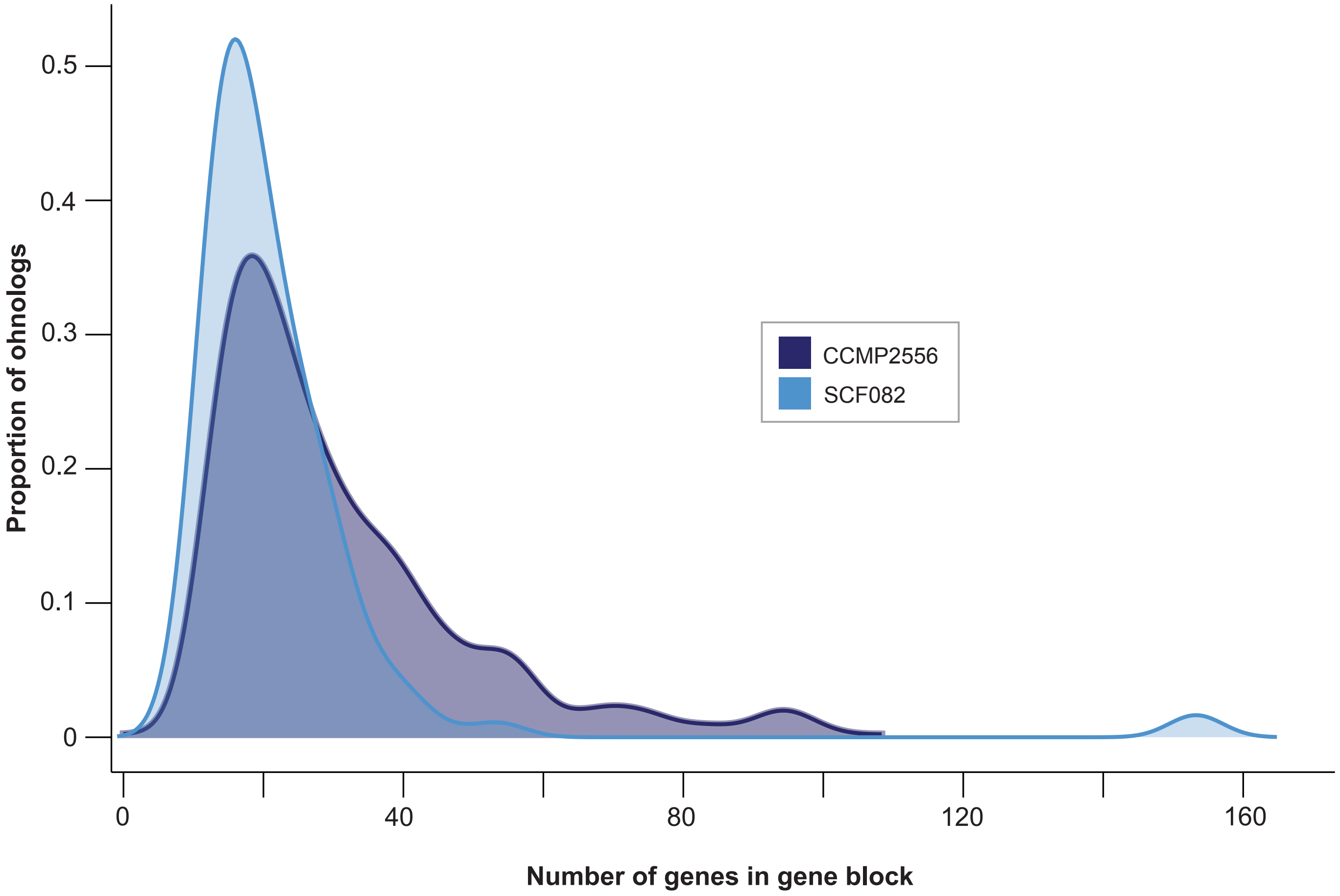


**Figure S2 |** Distribution of the number of genes per collinear gene block for the two *D. trenchii* isolates.

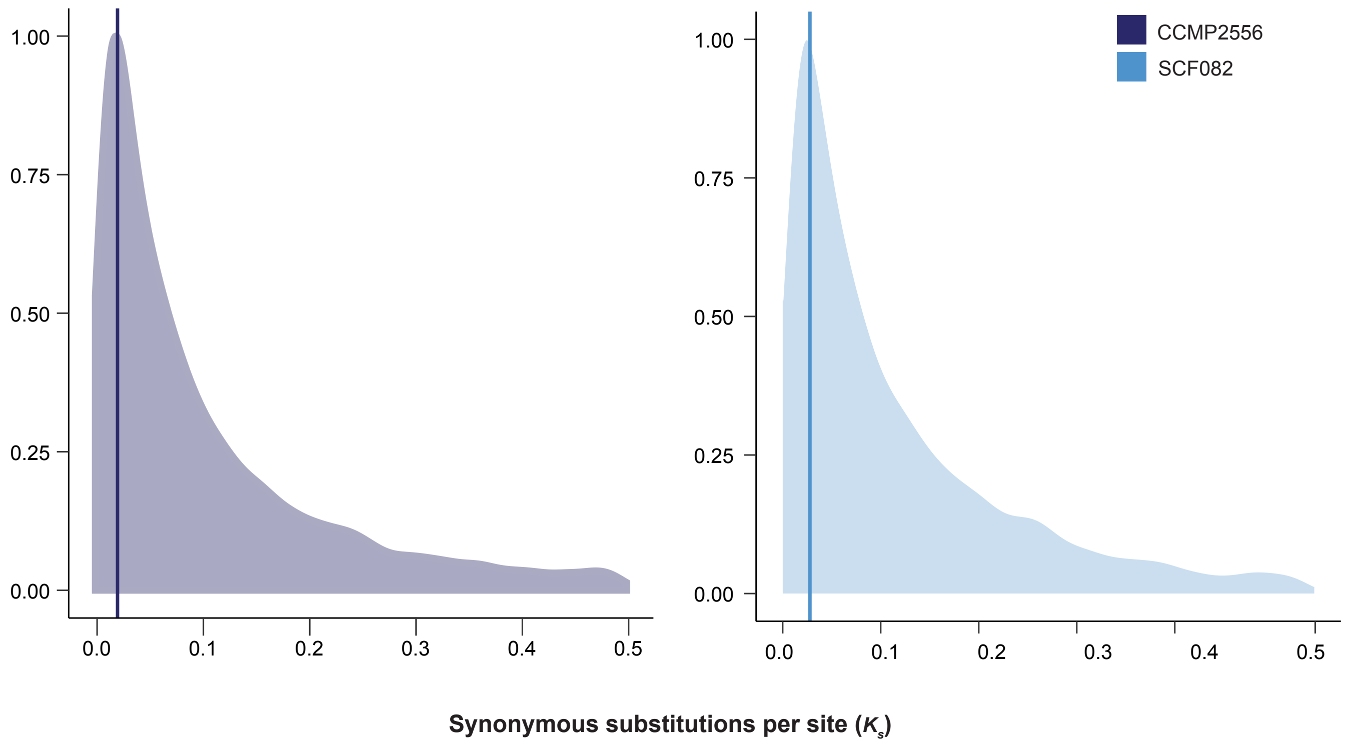


**Figure S3 |** Distribution of the number of synonymous susbstitutions per site (Ks) with lines indicating peaks for CCMP2556 and SCF082.

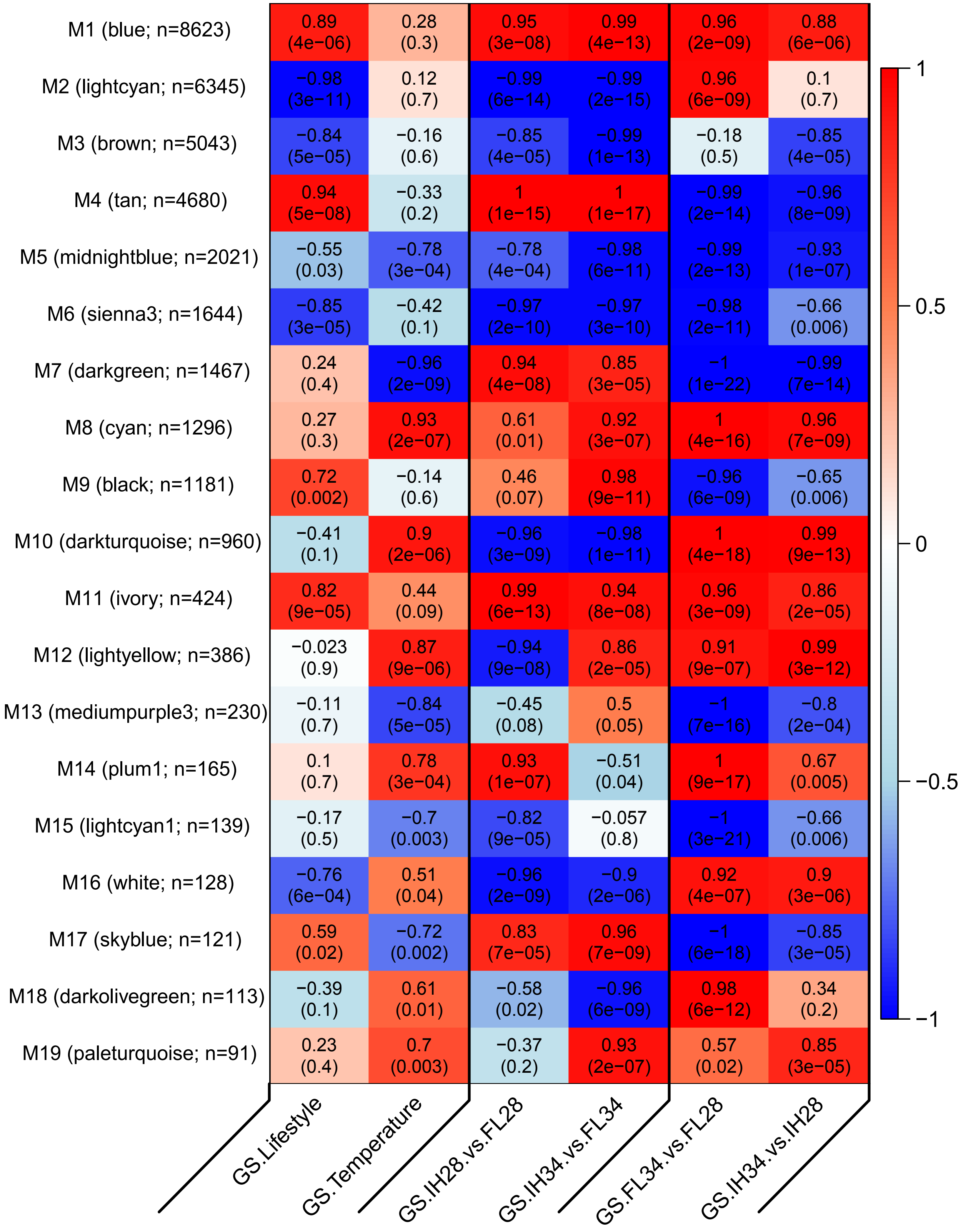


**Figure S4 |** Heatmap of the WGCNA co-expression module heatmaps with correlations and the p-values for their relationships to the RNA-seq treatments.

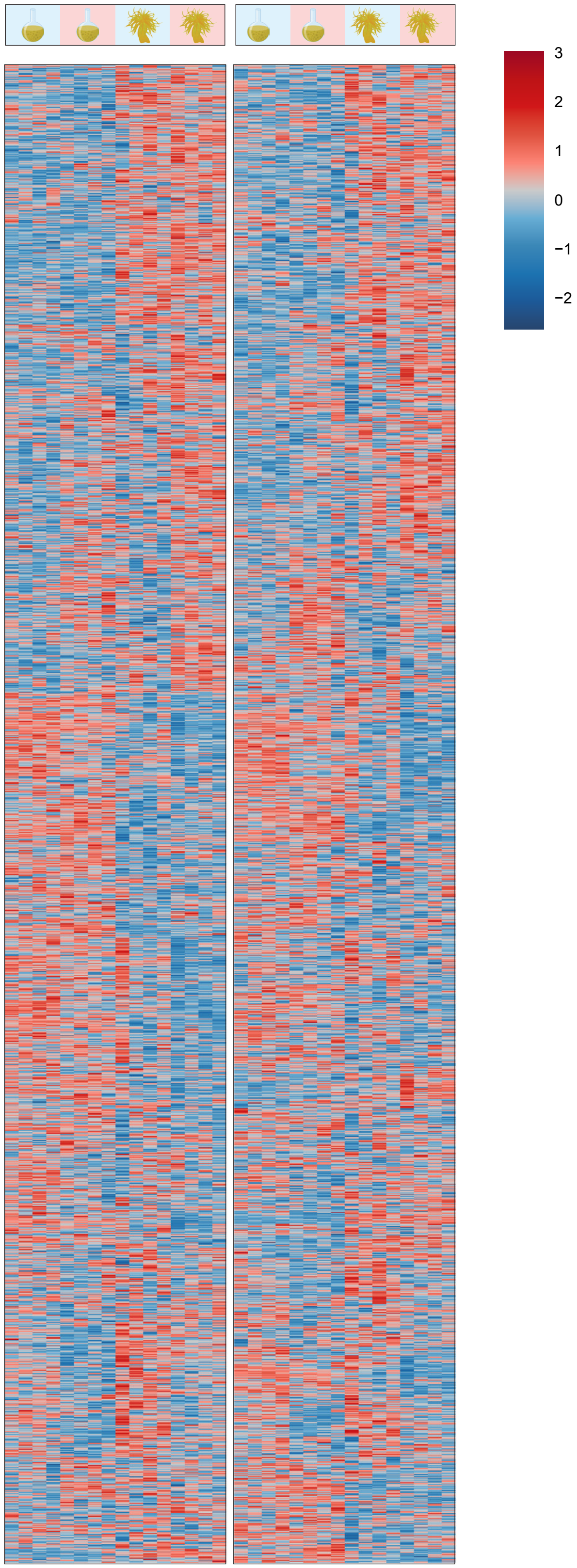


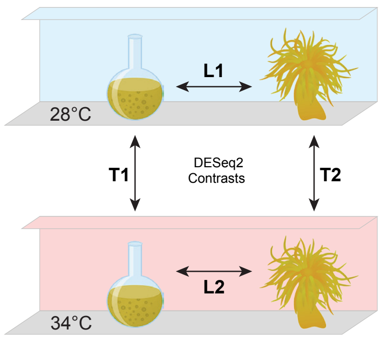


**Figure S5 |** Z-score normalized heatmap for Group 1 ohnolog pairs where neither ohnolog displayed any differential expression.

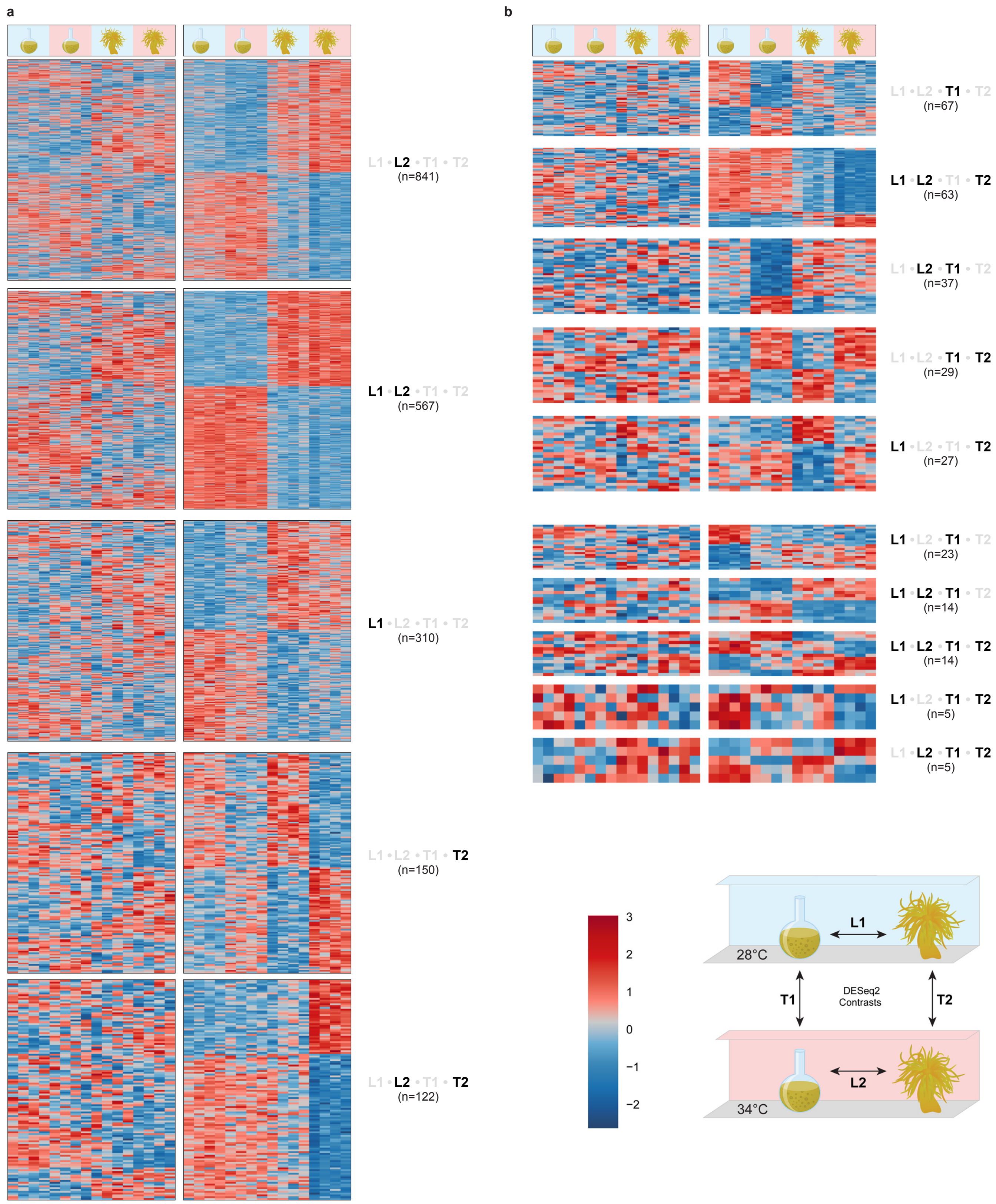


**Figure S6 |** Z-score normalized heatmap for Group 2 ohnolog pairs where only one of the two ohnologs displayed any differential expression. The differentially expressed ohnolog is represented on the right.

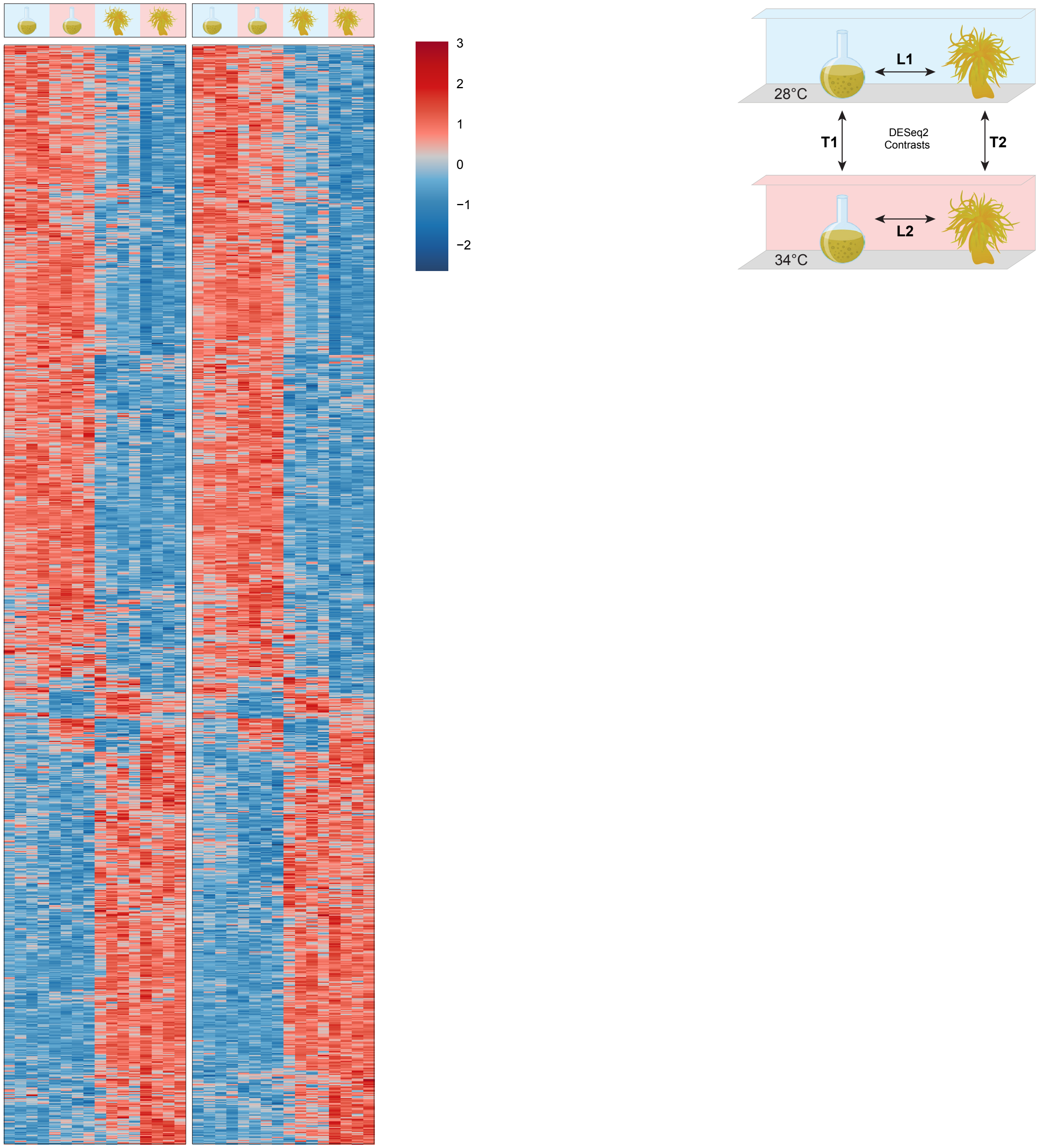


**Figure S7 |** Z-score normalized heatmap for Group 3 ohnolog pairs where both ohnologs displayed similar patterns of differential expression at the same time.

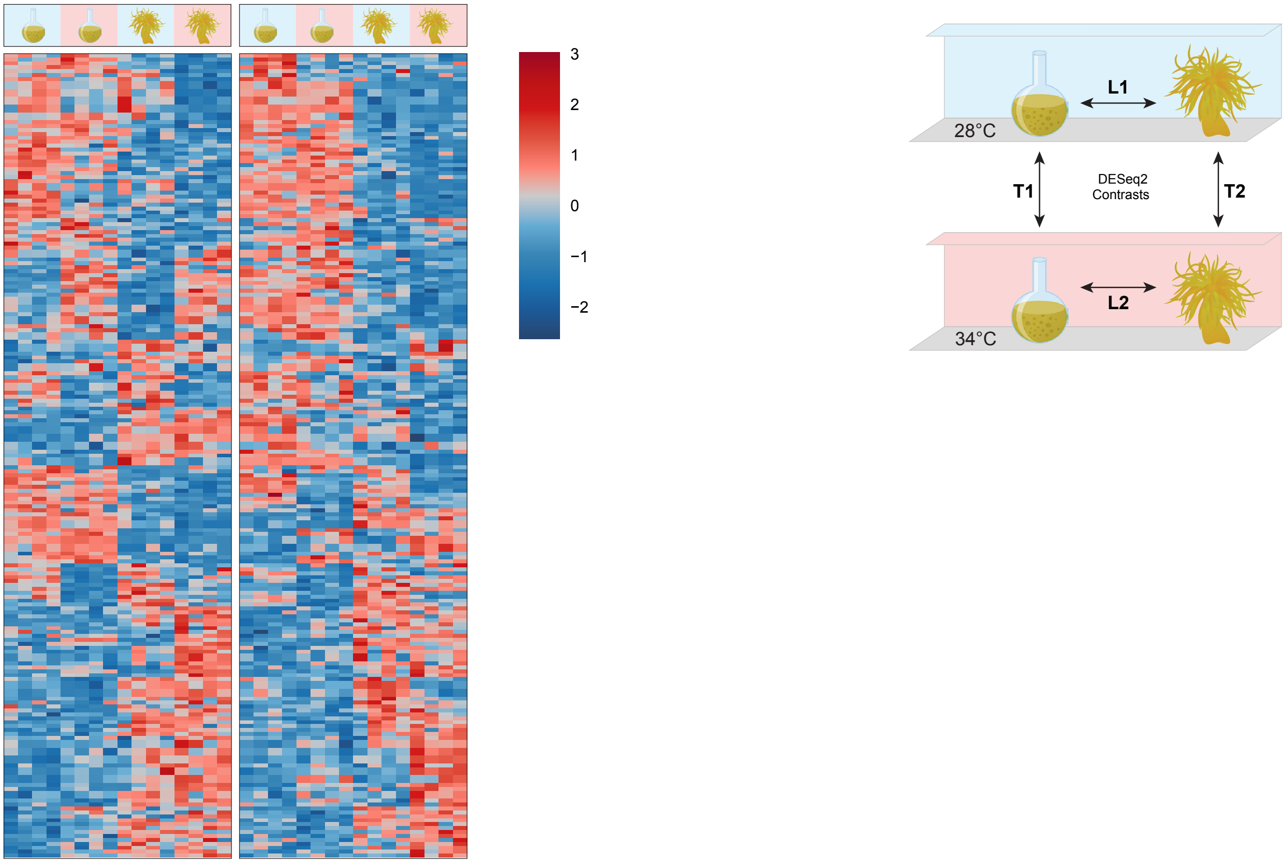


**Figure S8 |** Z-score normalized heatmap for Group 4 ohnolog pairs both ohnologs displayed differential expression but at different times.

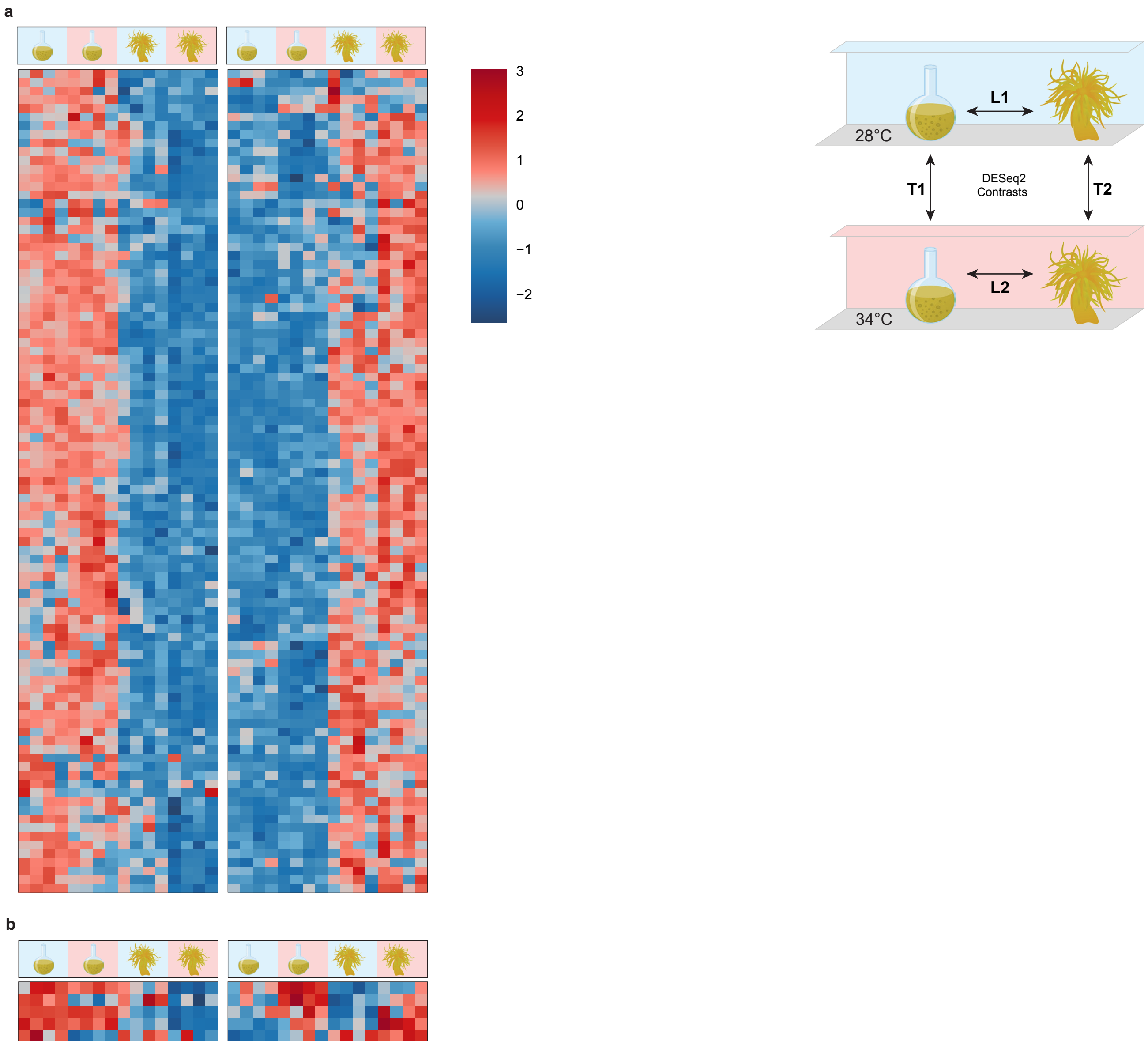


**Figure S9 |** Z-score normalized heatmap for Group 5 ohnolog pairs where both ohnologs displayed contrasting patterns of differential expression.

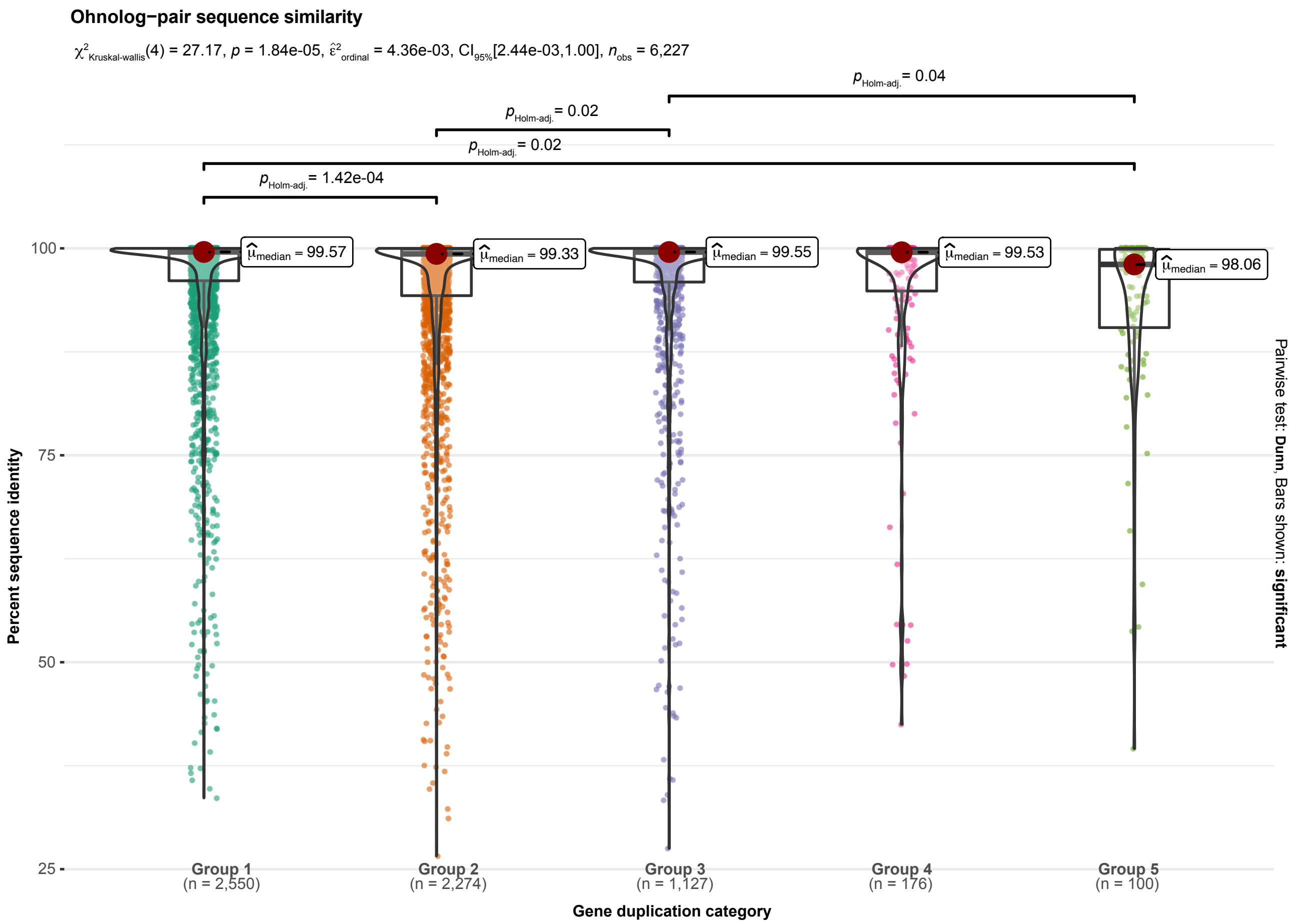


| Expression Group | Mean | Median |
|---|---|---|
| **Group 1** | 90.7 | 99.3 |
| **Group 2** | 82.6 | 97.6 |
| **Group 3** | 86.6 | 99.0 |
| **Group 4** | 90.2 | 99.1 |
| **Group 5** | 81.9 | 94.2 |

**Figure S10 |** Violin box plots of sequence similarity across the five groups with variable differential expression patterns.

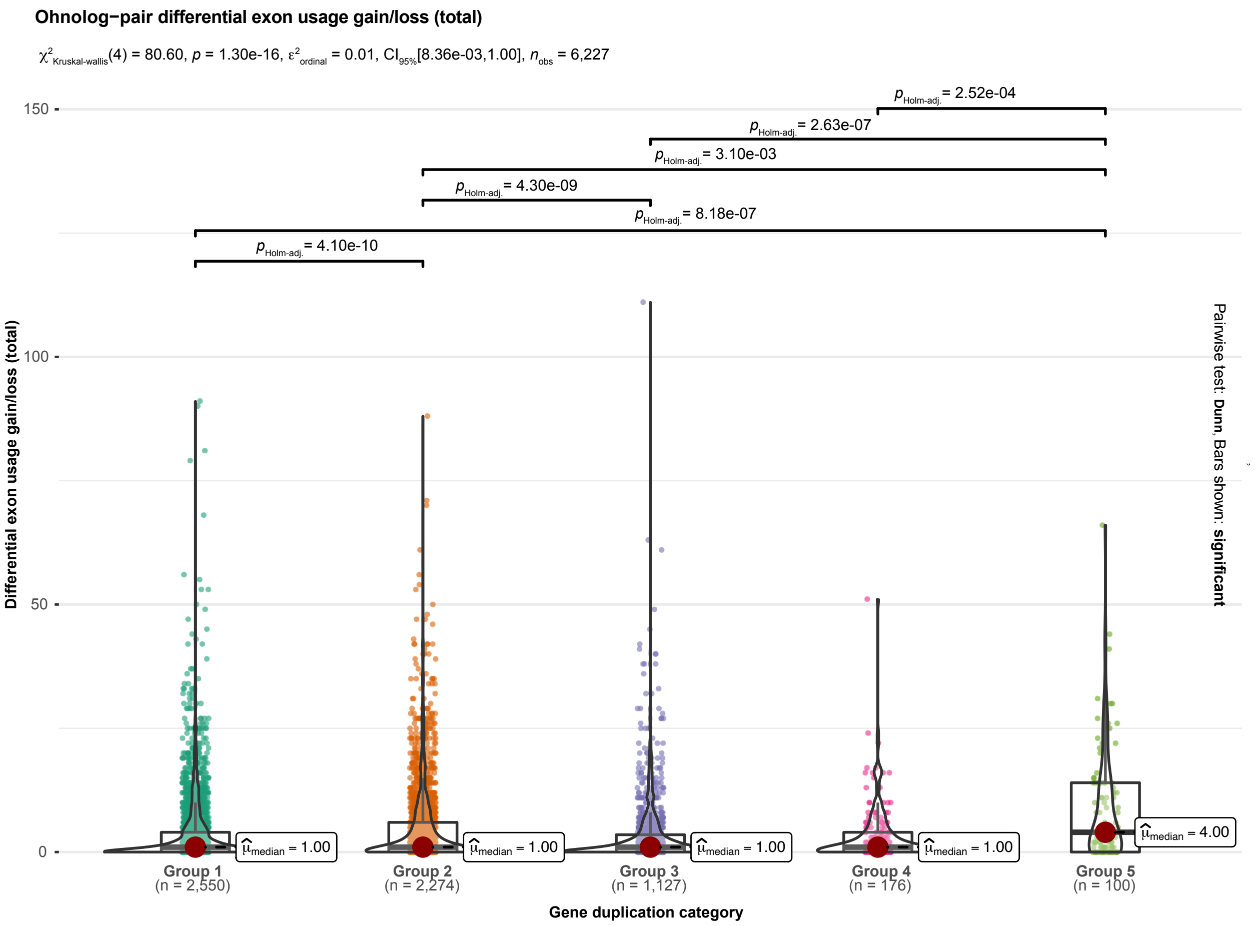


| Expression Group | Mean | Median |
|---|---|---|
| Group 1 | | |
| Group 2 | | |
| Group 3 | | |
| Group 4 | | |
| Group 5 | | |

**Figure S11 |** Violin box plots of the absolute difference in differential exon usage across the five groups with variable differential expression patterns.

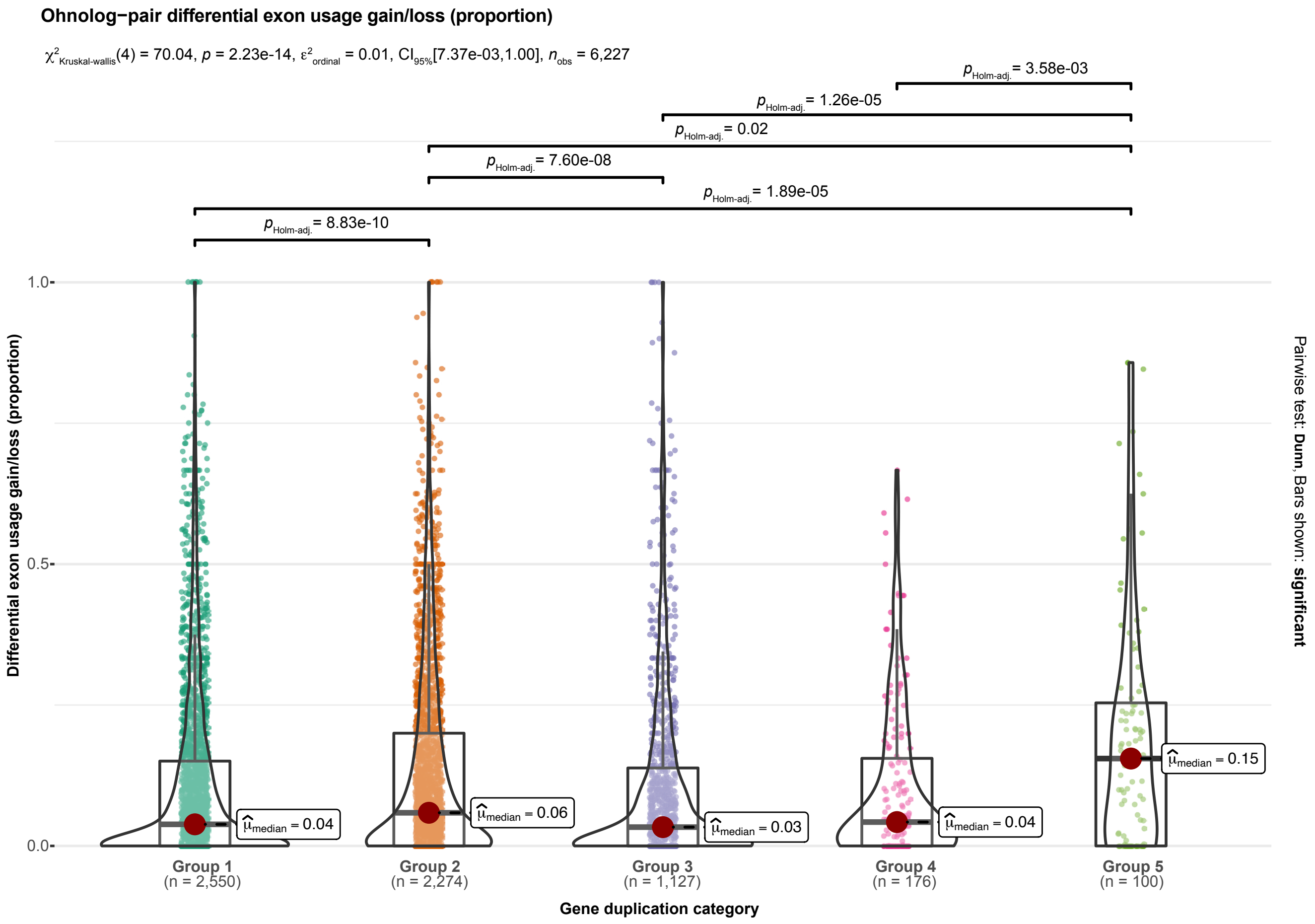


| Expression Group | Mean | Median |
|---|---|---|
| Group 1 | | |
| Group 2 | | |
| Group 3 | | |
| Group 4 | | |
| Group 5 | | |

**Figure S12 |** Violin box plots of the difference in differential exon usage, proportional to total number of exons, across the five groups with variable differential expression patterns.

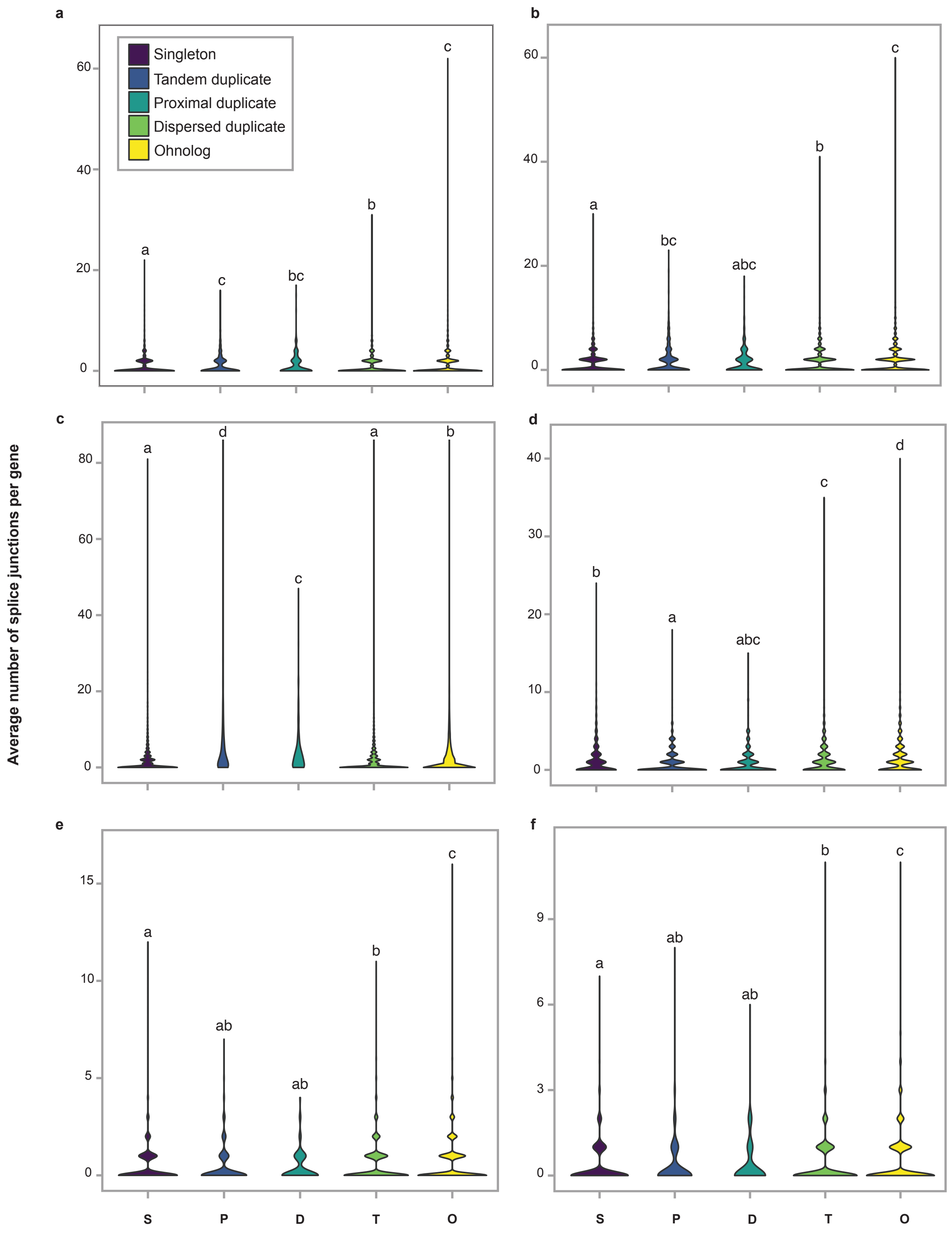


**Figure S13 |** Violin plots of the average number of splice junctions per gene across the gene du-plication categories. The average number of splice junctions per gene is shown for the splice junction types (A) alternate acceptor (B) alternate donor (C) alternate exon (D) retained intron (E) intron start and (F) intron end. Statistically different groups identified through a Kruskal-Willis post-hoc test are identified with letters above the plot.

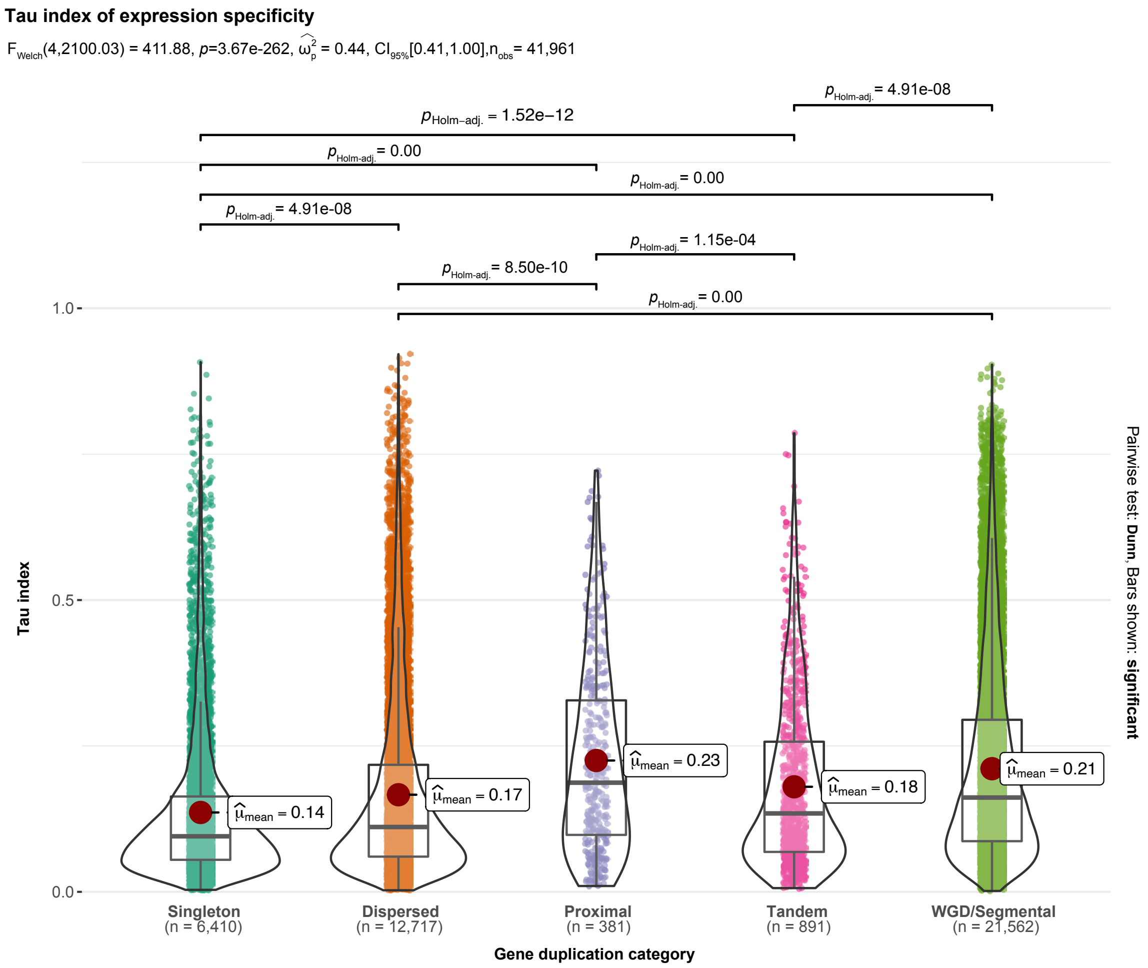


| Gene duplication type | Mean | Median |
|---|---|---|
| **Singleton** | 0.14 | 0.10 |
| **Dispersed** | 0.17 | 0.11 |
| **Proximal** | 0.23 | 0.19 |
| **Tandem** | 0.18 | 0.13 |
| **WGD/Segmental** | 0.21 | 0.16 |

**Figure S14 |** Violin box plots of tau expression specificy indices for the MCScanX categories.

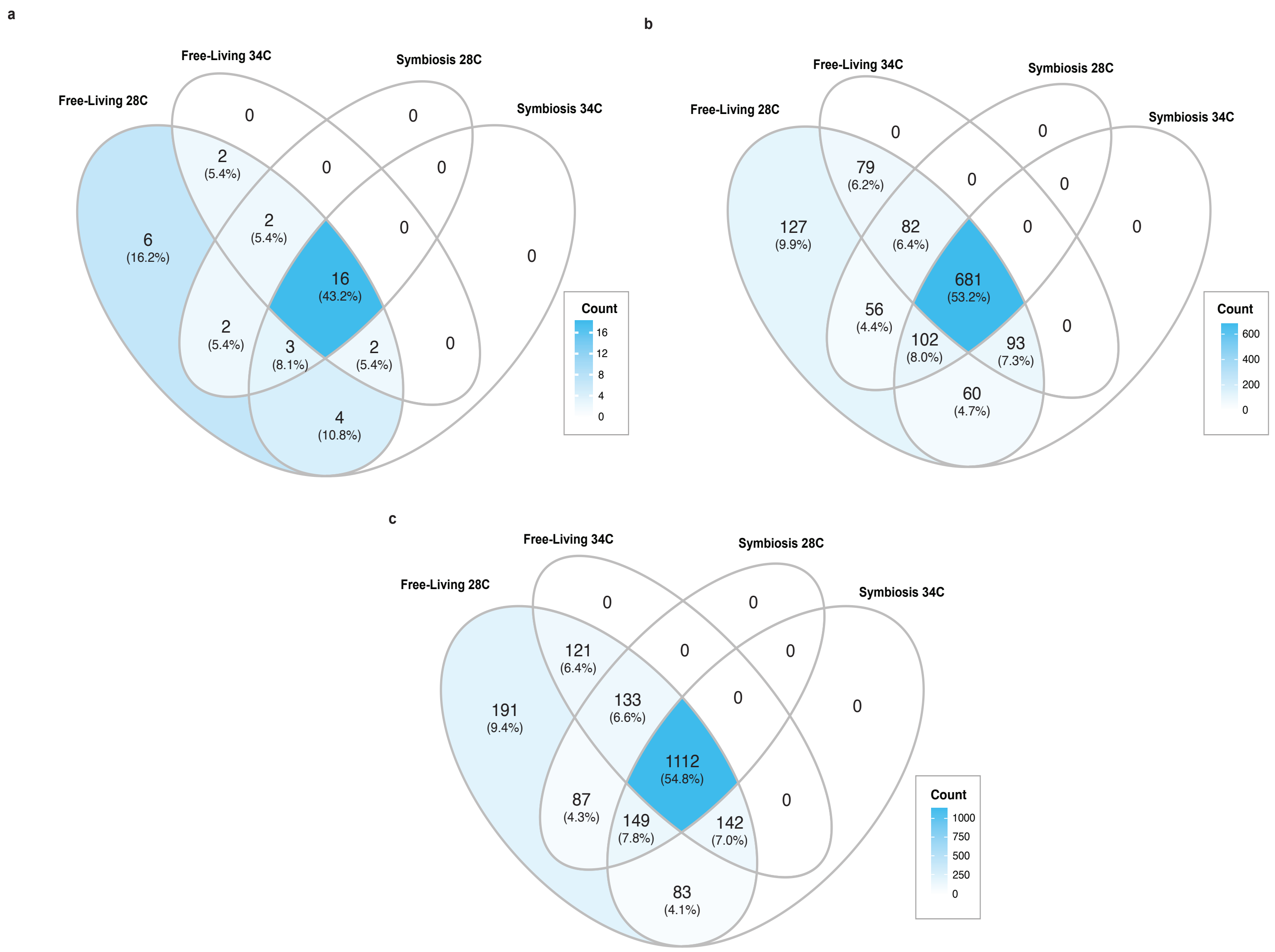


**Figure S15 |** Summary of consensus mRNA-edited sites in CCMP2556 across the four RNAseq treatments. The numbers represent the number of distinct edited sites for those that have a (a) high impact (ie. stop gained, stop lost, start lost) on gene function, (b) moderate (i.e. missense edit) impact on gene function, and (c) all consensus edited sites including synonymous changes.

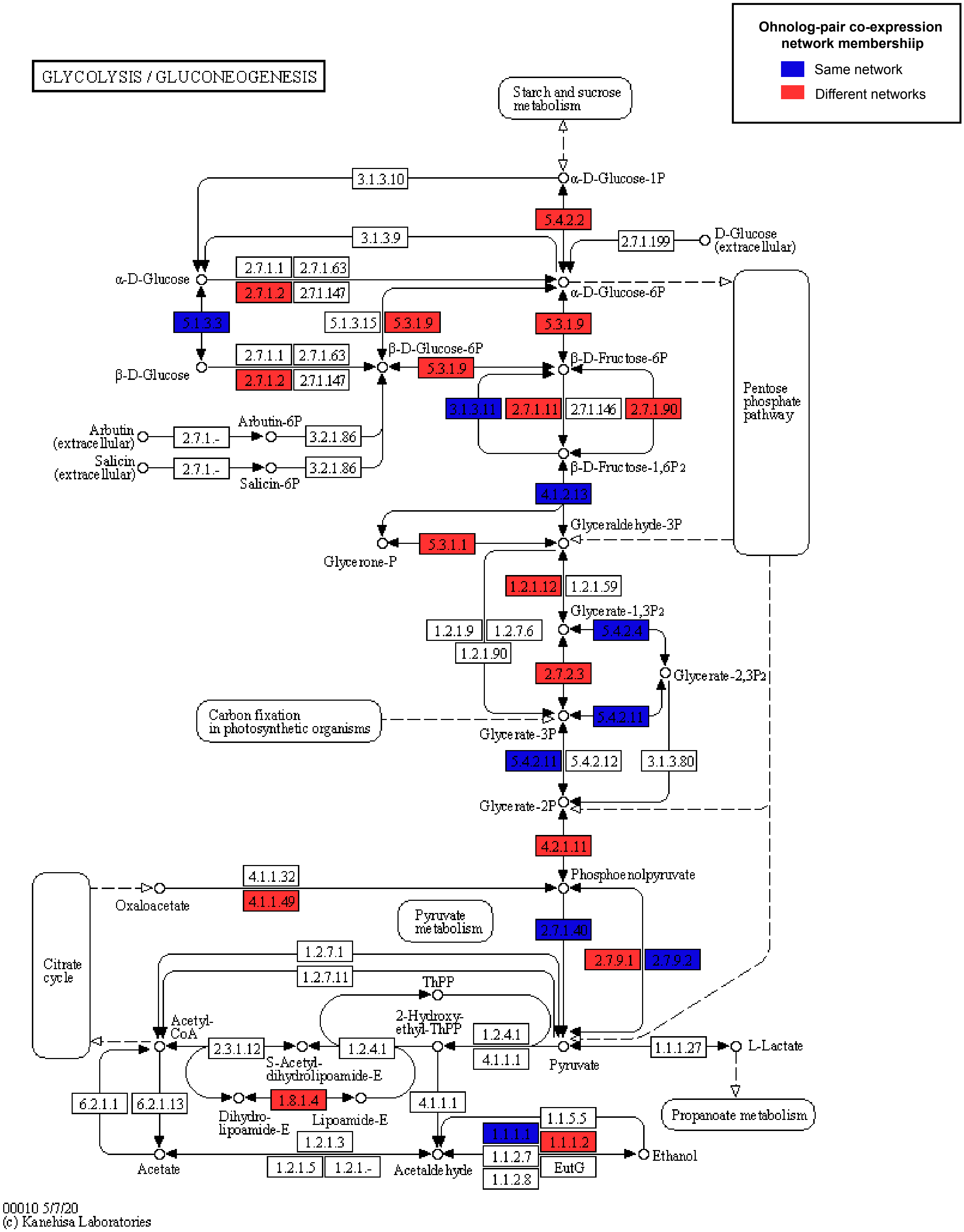


**Figure S16 |** KEGG pathway representation of the Glycolysis/Gluconeogenesis with ohnolog pair co-expression network membership indicated by the colors, with those in blue found in the same co-expression network and those in red in different co-expression networks.

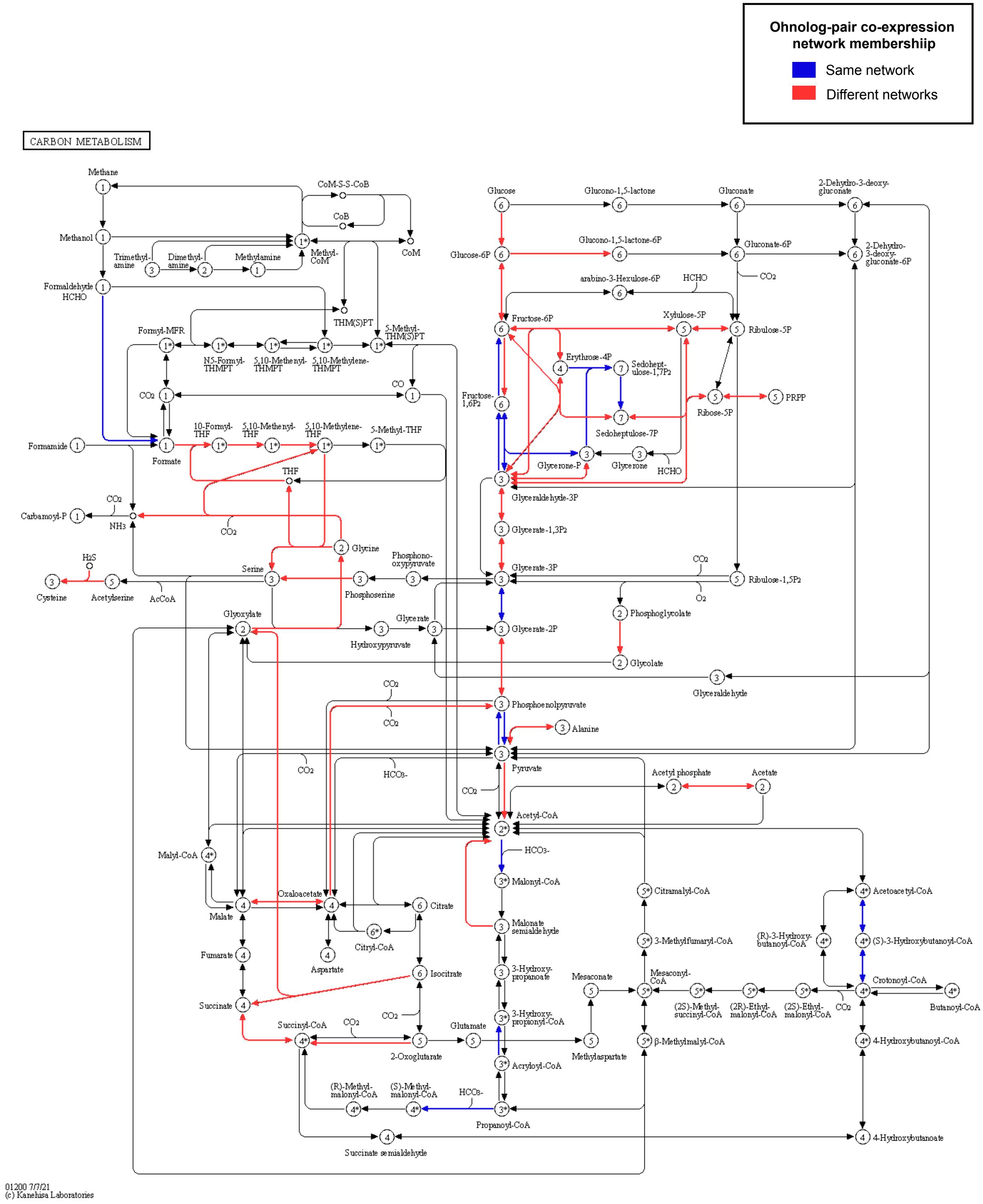


**Figure S17 |** KEGG pathway representation of Carbon Metabolism with ohnolog pair co-expression network membership indicated by the colors, with those in blue found in the same co-expression network and those in red in different co-expression networks.

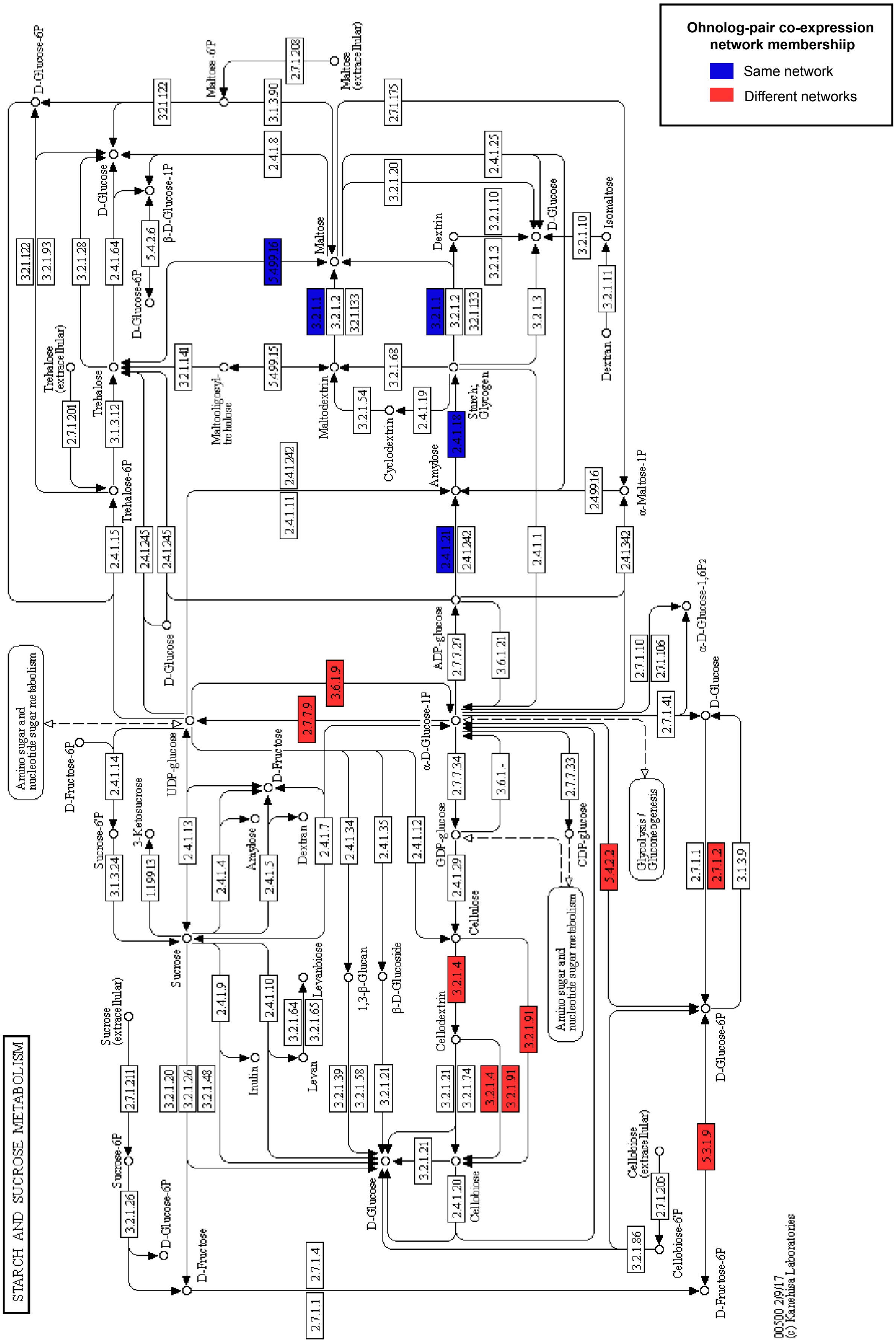


**Figure S18 |** KEGG pathway representation of Starch and Sucrose Metabolism with ohnolog pair co-expression network membership indicated by the colors, with those in blue found in the same co-expression network and those in red in different co-expression networks.

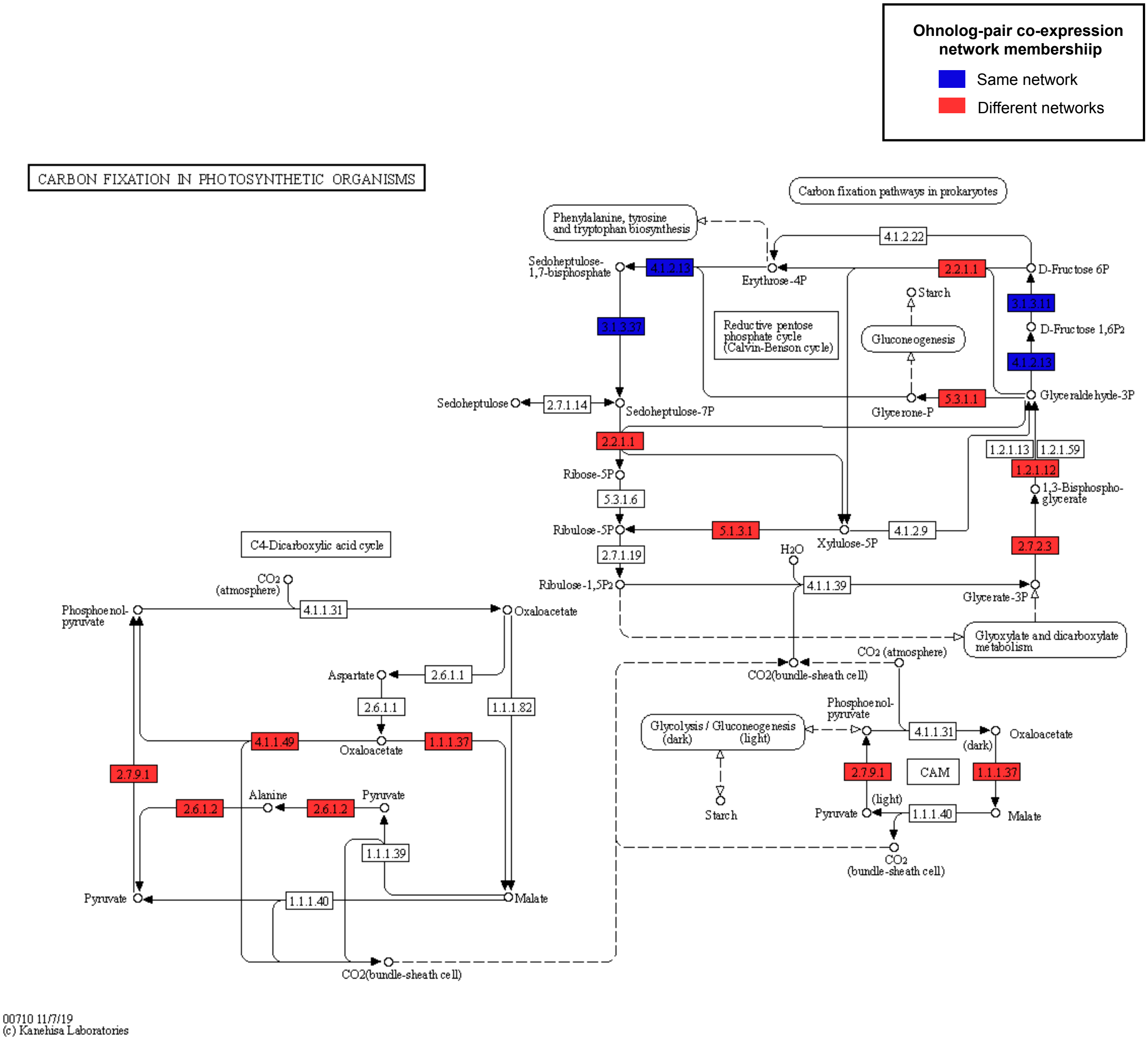


**Figure S19 |** KEGG pathway representation of Carbon Fixation in Photosynthetic Organisms with ohnolog pair co-expression network membership indicated by the colors, with those in blue found in the same co-expression network and those in red in different co-expression networks.

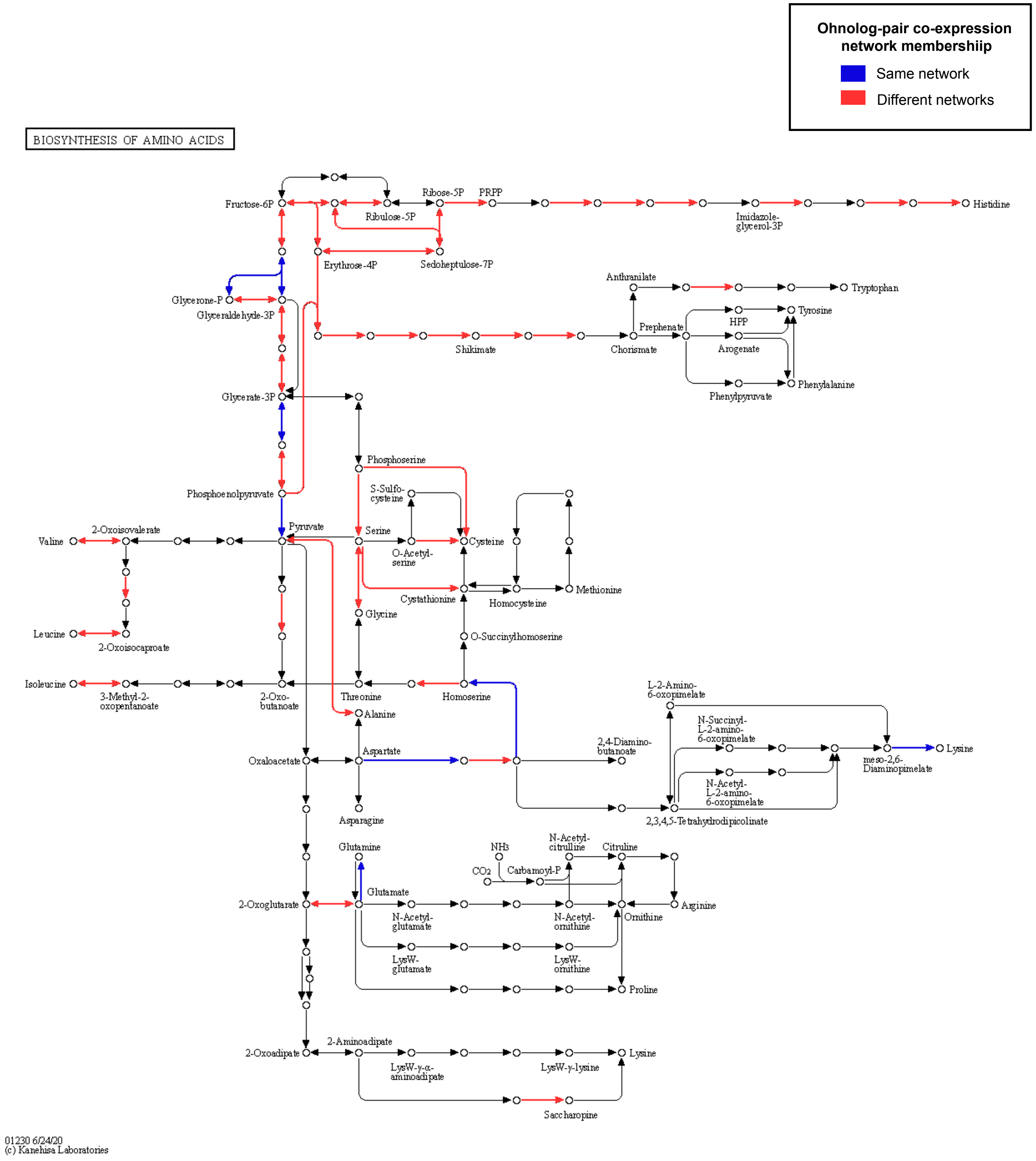


**Figure S20 |** KEGG pathway representation of Biosynthesis of Amino Acids with ohnolog pair co-expression network membership indicated by the colors, with those in blue found in the same co-expression network and those in red in different co-expression networks.

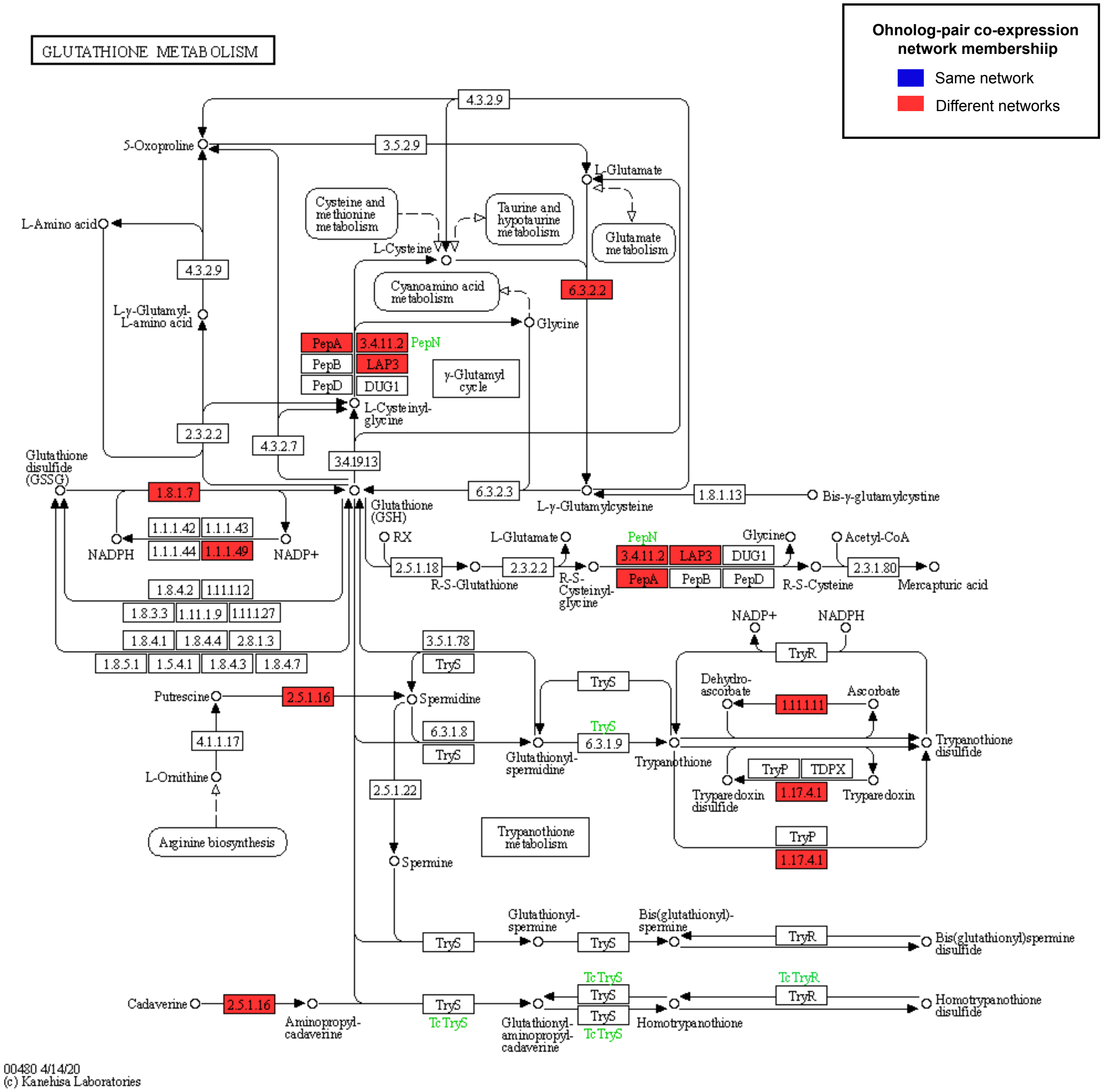


**Figure S21 |** KEGG pathway representation of Glutathione Metabolism with ohnolog pair co-expression network membership indicated by the colors, with those in blue found in the same co-ex-pression network and those in red in different co-expression networks.

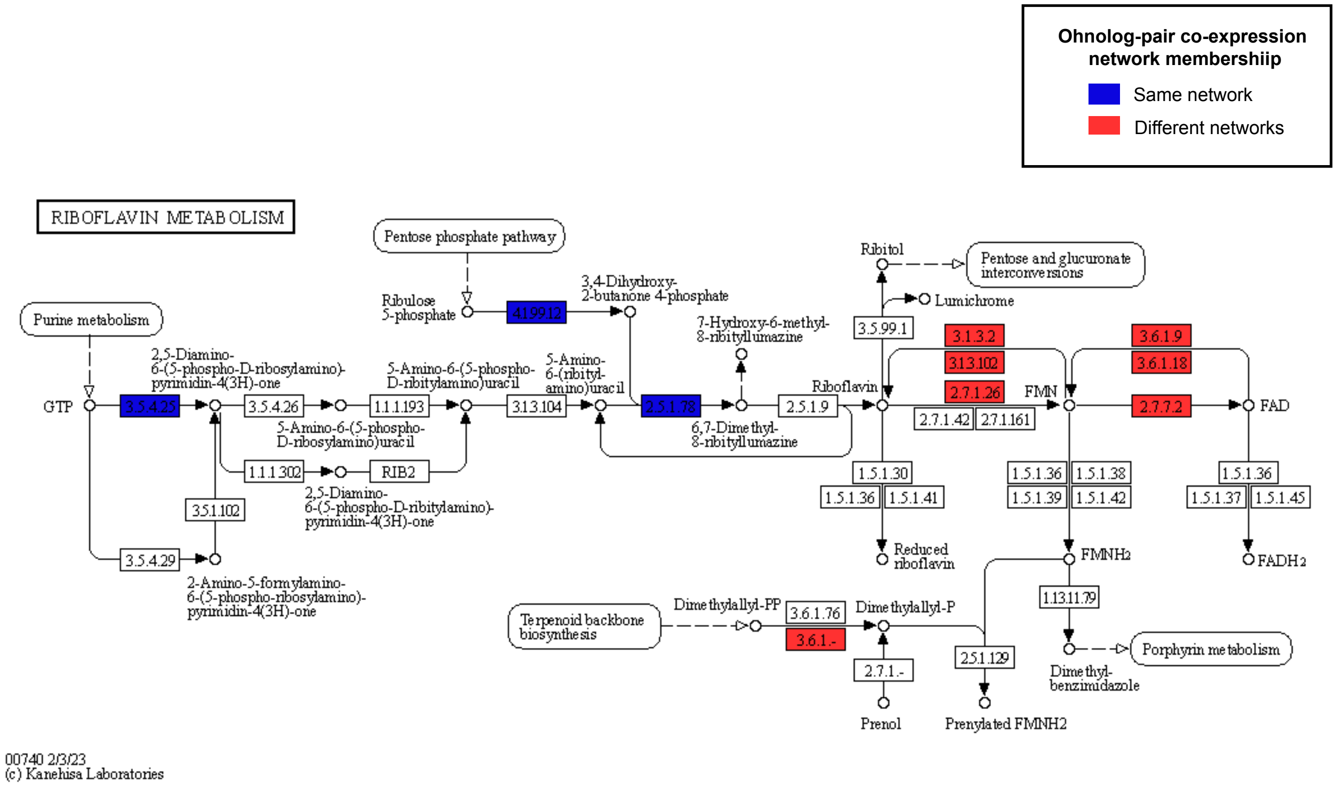


**Figure S22 |** KEGG pathway representation of Riboflavin Metabolism with ohnolog pair co-expression network membership indicated by the colors, with those in blue found in the same co-ex-pression network and those in red in different co-expression networks.

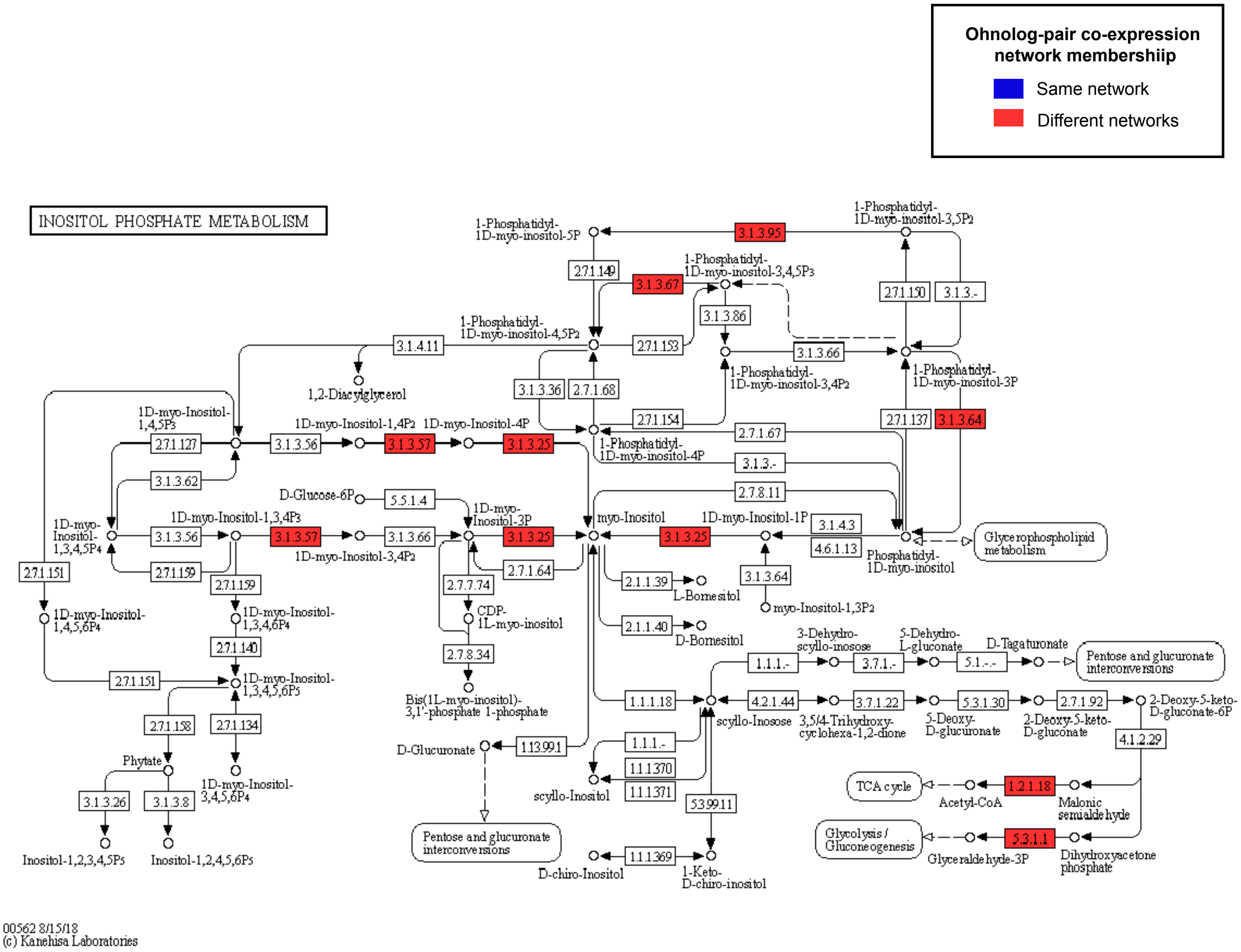


**Figure S23 |** KEGG pathway representation of Inositol Phosphate Metabolism with ohnolog pair co-expression network membership indicated by the colors, with those in blue found in the same co-expression network and those in red in different co-expression networks.

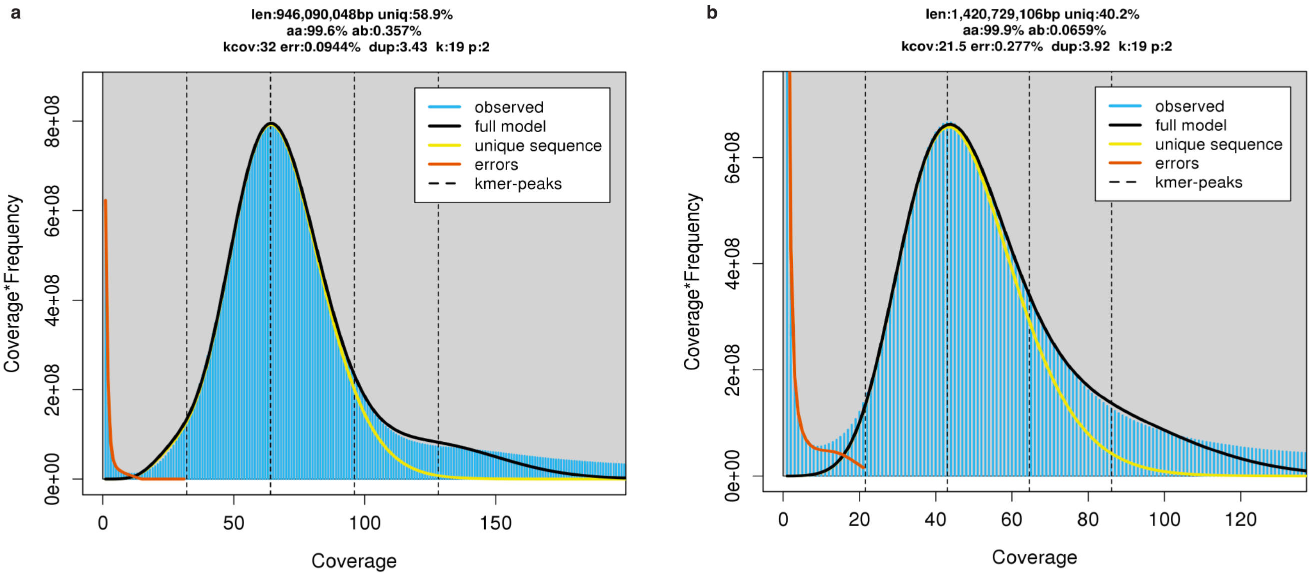


**Figure S24 |** *Durusdinium trenchii* GenomeScope2 plots.

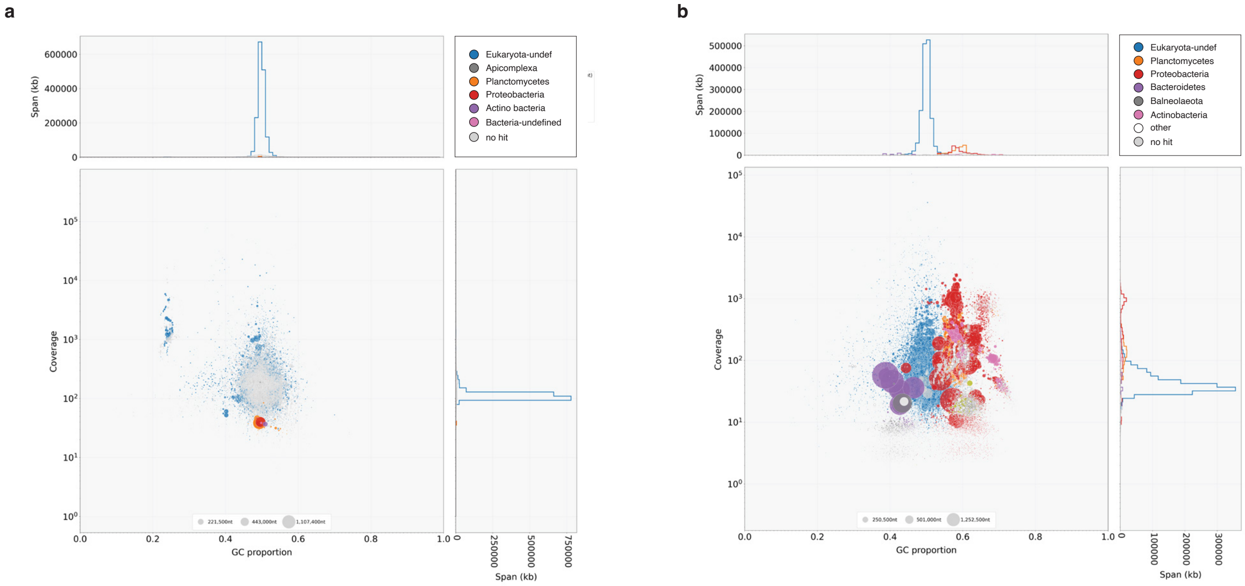


**Figure S25 |** Blobtools taxon-annotated plot for *D. trenchii* (a) CCMP2556 and (b) SCF082.
